# The CCCH-type zinc-finger *Pf*MD3 promotes translation for malaria parasite transmission

**DOI:** 10.64898/2026.05.27.726058

**Authors:** Riëtte van Biljon, Suyash Bhatnagar, Rajat Kumar, Megan Gliozzi, Guoyue Xu, Abhai Tripathi, Arnab Ashrafi, Kristian Swearingen, Photini Sinnis, Manuel Llinás, Heather J. Painter

**Affiliations:** Division of Bacterial, Parasitic, and Allergenic Products, Office of Vaccines Research and Review, Center for Biologics Evaluations and Research, Food and Drug Administration, Silver Spring, MD, USA; Department of Biochemistry & Molecular Biology and the Huck Center for Malaria Research, Pennsylvania State University, University Park, PA, USA; Department of Chemistry, Pennsylvania State University, University Park, PA, USA; Johns Hopkins Malaria Research Institute and Department of Molecular Microbiology & Immunology, Baltimore, MD, USA; Institute for Systems Biology, Seattle, WA, US

## Abstract

Sexual maturation and fertility of the malaria causing parasite, *Plasmodium* spp., is critical to transmission between hosts. This developmental process involves an intricate gene expression regulatory network that drives gametocyte commitment, development, and the production of male and female gametes. Here, we characterized the role of the CCCH-type zinc-finger protein *Plasmodium falciparum* Male Development 3 (*Pf*MD3) in the formation of gametes. Genetic disruption of *Pf*MD3 revealed a male fertility defect that decreased infectivity to mosquitoes. Molecular characterization of *Pf*MD3 revealed its binding to mRNA transcripts associated with male gamete development through a specific RNA sequence. Gametocyte proteins encoded by these *Pf*MD3-bound transcripts decreased significantly in *Pf*MD3 knockout parasites, suggesting that *Pf*MD3 impacts translation. In addition to RNA-binding, *Pf*MD3 also interacts with proteins involved in mRNA processing and nuclear export that influence translational regulation. This study establishes *Pf*MD3 as an RNA-binding protein that impacts lineage-specific translation and male gamete fertility.

## Introduction

Malaria is a disease that has plagued humankind for millennia and continues to claim over half a million lives each year^1^. The causative pathogen of malaria is the *Plasmodium* parasite which, through a sophisticated balance of infection and replication, shuttles between the female anopheline mosquito host and the human host to complete its lifecycle^2^. During human infection, *Plasmodium* parasites grow and replicate asexually within red blood cells (RBCs). Cyclical invasion and rupture of RBCs (every 48 hours for *P. falciparum*) results in the clinical symptoms of malaria. In each cycle, a subset of blood-stage parasites converts to sexual stage gametocytes, which are the only parasite form that is infectious to female *Anopheles* mosquitoes during a blood meal from an infected host^3,4^. Thus, targeting processes or proteins essential to the formation of gametocytes or the fertilization of gametes presents an effective therapeutic intervention strategy to block parasite transmission^5^.

Much effort has been dedicated to understanding the molecular and cellular factors behind the phenotypic and metabolic changes that occur during gametocyte commitment. Numerous whole genome studies have characterized the gametocyte transcriptome and its potential regulators demonstrating that the parasite rewires its gene expression program to switch from replicating asexual blood-stage development to terminally differentiated sexual cellular forms^6–24^. Furthermore, gametocyte development entails morphological differentiation into male and female sexual precursors to ensure production of micro- and macrogametes in the mosquito. Therefore, sex-specific regulatory processes are required, although it remains an open question as to when and how sexual fate is determined^3,25,26^.

Despite the importance of DNA-binding proteins (reviewed in^27^) and epigenetic factors^13,28–30^ in regulating the transition of *P. falciparum* parasites through distinct lifecycle stages, the number of DNA binding proteins encoded in the genome is low^31,32^.

On the other hand, hundreds of RNA-binding proteins have been predicted in the parasite genome^33–37^ suggesting that gene expression in *Plasmodium* parasites is regulated transcriptionally, post-transcriptionally^6,38–40^, and at the level of translation^41–52^. Historically, studies have shown that RNA metabolism in *P. falciparum* slows from stage III through stage V of gametocyte development^53,54^, suggesting a reliance on post-transcriptional regulation for the final stages of maturation and fertilization. Despite the abundance of RNA-binding proteins and evidence supporting RNA-centric mechanisms of gene regulation during the long maturation of *P. falciparum* gametocytes, only a handful of proteins that possess nucleic acid interacting domains have been studied in-depth to date^55–58^. Specifically, Pumilio family (PUF) proteins^46,48,59^, an OST-HTH/LOTUS domain containing protein (*Pf*MD1)^60^, Male Development Protein 5 (MD5, PF3D7_0414500)^61^ also known as RBPm1 in *P. yoelii*^40^, the RNA helicase <u>D</u>evelopment <u>o</u>f <u>Z</u>ygote Inhibited (DOZI)^49^, and an Sm-like homolog of worm <u>C</u>AR-I and fly <u>T</u>railer <u>H</u>itch (CITH)^44^ have all been shown to be essential for sexual maturation and fertilization. Interestingly, the majority of the phenotypically characterized RNA-binding proteins play a role in the production of fertile female gametocytes^43–48,62,63^, likely influencing the timed translation of target transcripts in gamete and sporozoite mosquito stages^42,43^.

The *P. falciparum* genome encodes a large understudied family of ∼170 zinc finger (ZnF) domain-containing proteins, which are categorized based upon the arrangement of the cysteine and histidine residues within the ZnF domain^64,65^. The most highly represented ZnF domain arrangements are the CCCH-, RING-, PhD- and C2H2-types^65,66^. In higher eukaryotes, CCCH-type ZnF domain-containing proteins have been shown to perform a wide range of molecular functions including mRNA splicing, polyadenylation, decay, and promoting translation^67,68^, many of which rely on both sequence-specific RNA binding and a range of protein-protein interactions. Of the CCCH-type ZnF proteins encoded by the *P. falciparum* genome, a few have been experimentally linked to roles in mRNA regulation. For example, *Pf*CZIF1 and 2 were shown to regulate schizogony through controlling gene expression^69^. In gametocytes, several proteins with ZnF domains (ZNF4, RNF1 and GD1) have been shown to regulate maturation and fertility^70–72^. Similarly, PF3D7_0315600 (*Pf*MD3)^61^, was found to be important for male gametocyte development through a network of interactions with regulators of gametocytogenesis, including other RNA-binding proteins^73^, but its specific role in mRNA regulation remains undefined.

In this study, we provide an in-depth characterization of *Pf*MD3 as an RNA-binding protein that regulates the translation of gametocyte proteins which are essential for the transmission of *P. falciparum*. We demonstrate that *Pf*MD3 is preferentially expressed in male gametocytes and plays a critical role in male gametogenesis. Genetic disruption of *Pf*MD3 resulted in a pronounced decrease in mosquito transmission and in a reduction of P230p positive male gametocytes. By analyzing the transcriptome and proteome of the *Pf*MD3 knockout parasite line, we found a significant decrease in the abundance of male-associated transcripts and proteins. These findings are corroborated by single cell RNA-sequencing data that demonstrate a reduction in the male lineage of gametocytes lacking *Pf*MD3. We show that *Pf*MD3 is an RNA binding protein that directly binds to a sequence-specific motif in mRNA transcripts that are necessary for male gamete development. Protein-protein interactions with *Pf*MD3 identify an enrichment for RNP biogenesis proteins and components of the translational machinery. Through a multi-omics integrative analysis of the datasets produced herein, we demonstrate that *Pf*MD3 binds specific mRNA targets to promote their translation to produce fertile male gametocytes.

## Results

### *Pf*MD3 is a CCCH-type zinc finger protein expressed throughout intraerythrocytic development

*Pf*MD3 is a 485 amino acid protein with a well conserved CCCH-type zinc finger domain (Figure 1A; aa 163-185) and a predicted lysine-proline rich ribosome-binding receptor domain: RRD_K/P (Figure 1A; aa 260-295) found only in species of the subgenus *Laverania* (Figure 1B). The *Pf*MD3 protein is also predicted to contain two nucleolar localization signals, one of which overlaps with the RRD_K/P domain^74,75^ (Figure 1A; Document S1). From previous studies, the *pfmd3* transcript peaks in abundance in early gametocyte stages^7,73^. To confirm the timing and level of *Pf*MD3 protein expression relative to its transcript abundance, we integrated a 3xHA tag at the 3’-end of the *Pf*MD3 locus (Figures S1A, B) in *P. falciparum* strain NF54 Patient Line E (NF54e; MRA-1000).

**Figure 1.**
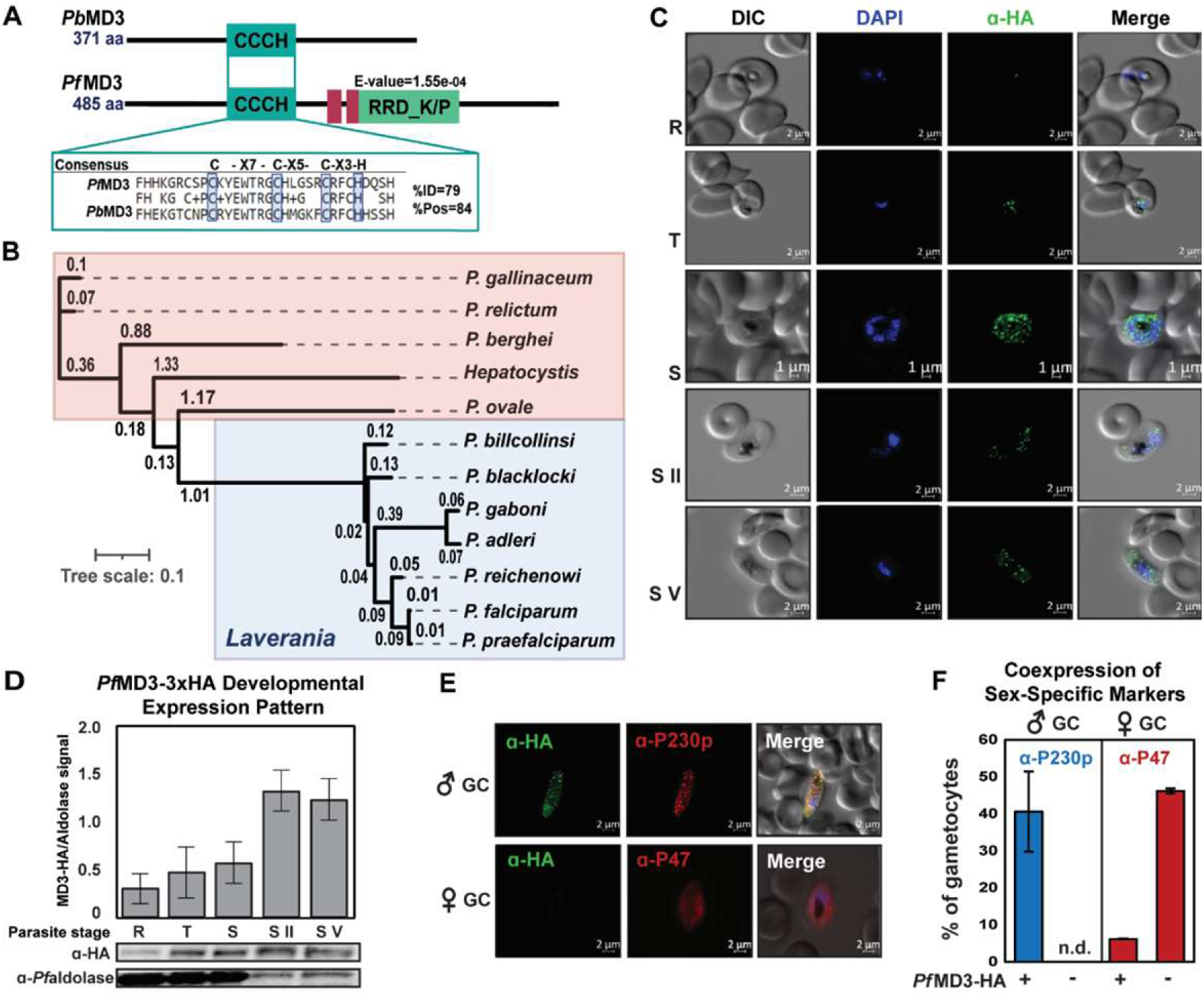
*Pf*MD3 evolutionary divergence and expression in human blood stage malaria parasite development. (**A**) Schematic representation of the protein domain architecture of both the *Pb*MD3 and *Pf*MD3 proteins. CCCH = CCCH-type zinc finger domain (IPR000571), RRD_K/P = Lysine-Proline rich ribosome-binding receptor domain (pfam05104, E-value = 1.55e-04), Magenta blocks indicate nucleolar localization signals (Document S1). Enlarged region shows the amino acid sequence of the CCCH-type domain with the consensus sequence between species indicated within the blue blocks. (**B**) Phylogenetic tree of *Pf*MD3 orthologs scaled by similarity noting the *Laveranian* species indicated in blue box and non-*Laveranian* orthologs in the red box. (**C**) Expression and cellular localization of *Pf*MD3-3xHA throughout the intraerythrocytic lifecycle assessed by immunofluorescence microscopy. Fixed *in vitro* cultured parasites were collected at R = ring, T = trophozoite, S = Schizont, S II = Gametocyte Stage II (day 5 post-induction), and S V = Gametocyte Stage V (day 11 post-induction), then stained with an anti-HA primary antibody and Alexa-488 coupled secondary antibody (*Pf*MD3, green), and DAPI (blue) to show location of the cell nucleus. Representative images for each developmental stage are shown. (**D**) Western blot capturing the expression of *Pf*MD3-3xHA across the blood stages probed with an anti-HA antibody and an anti-*Pf*aldolase loading control, representative of n = 3 blots and with image signal quantified in the adjacent graph using ImageJ software. Plotted are the mean ± SEM. (**E**) Representative images of male (P230p and DAPI positive) and female (P47 positive and DAPI positive) Stage V gametocytes. (**F**) Quantification of sex-specific *Pf*MD3 expression in the *Pf*MD3-HA line using immunofluorescence microscopy (n = 2) by co-staining Stage V gametocytes with a male-specific anti-P230p (red)^87^ or female-specific anti-P47 (red), anti-HA (*Pf*MD3, green) and DAPI (blue). Results are represented as average number of gametocytes with the noted staining patterns out of >30 gametocytes (n = 2) counted ± SEM where the blue bar represents males, and red represents females. n.d. = not detected.

Using immunofluorescent microscopy and western blots, we found that *Pf*MD3-3xHA is expressed throughout the asexual and gametocyte blood stages (Figures 1C, D, S1C) with expression peaking in early gametocytes (Stage II) (Figures 1D, S1C). *Pf*MD3-3xHA is localized to discrete puncta within the nucleus and cytoplasm (Figure 1C). Using co-staining with antibodies to male (P230p) or female (P47) specific markers, we observed that all male gametocytes (P230p positive, Stage IV-V) expressed *Pf*MD3-3xHA while only a small percentage of female (P47 positive) cells expressed *Pf*MD3-3xHA (Figures 1E and F). Taken together, these results demonstrate that *Pf*MD3 is expressed throughout both the asexual and sexual blood stages but is most highly expressed in male gametocytes.

### *Pf*MD3 is a critical factor for producing fertile microgametes

To determine the relevance of *Pf*MD3 to parasite development, we used CRISPR/Cas9 to target and delete the first two exons of the *pf3d7_0315600* coding sequence in wild-type *P. falciparum* NF54e (Figure S2A) resulting in transgenic Δ*Pf*MD3 parasites (Figure 2A). We confirmed the genetic disruption in two clones, Δ*Pf*MD3^1.1^ and Δ*Pf*MD3^1.2^ by genotyping PCR (Figure S2B) and whole genome sequencing (Figures S2C) which revealed a loss in *pfmd3* mRNA abundance in gametocytes by RNA sequencing from Δ*Pf*MD3^1.1^ (Figure S2D; Table S1). We also confirmed that no unique peptides corresponding to *Pf*MD3 were detected by mass spectrometry in the Δ*Pf*MD3^1.1^ parasites, while the protein was well-detected in NF54e wild-type parasites (Figure S2E). Surprisingly, genetic disruption of *Pf*MD3 resulted in a significant increase in the asexual multiplication rate (Figure 2B) due to a slight increase in the percent parasitemia of both Δ*Pf*MD3 clones over 72 hours of growth compared to the NF54e parent.

**Figure 2.**
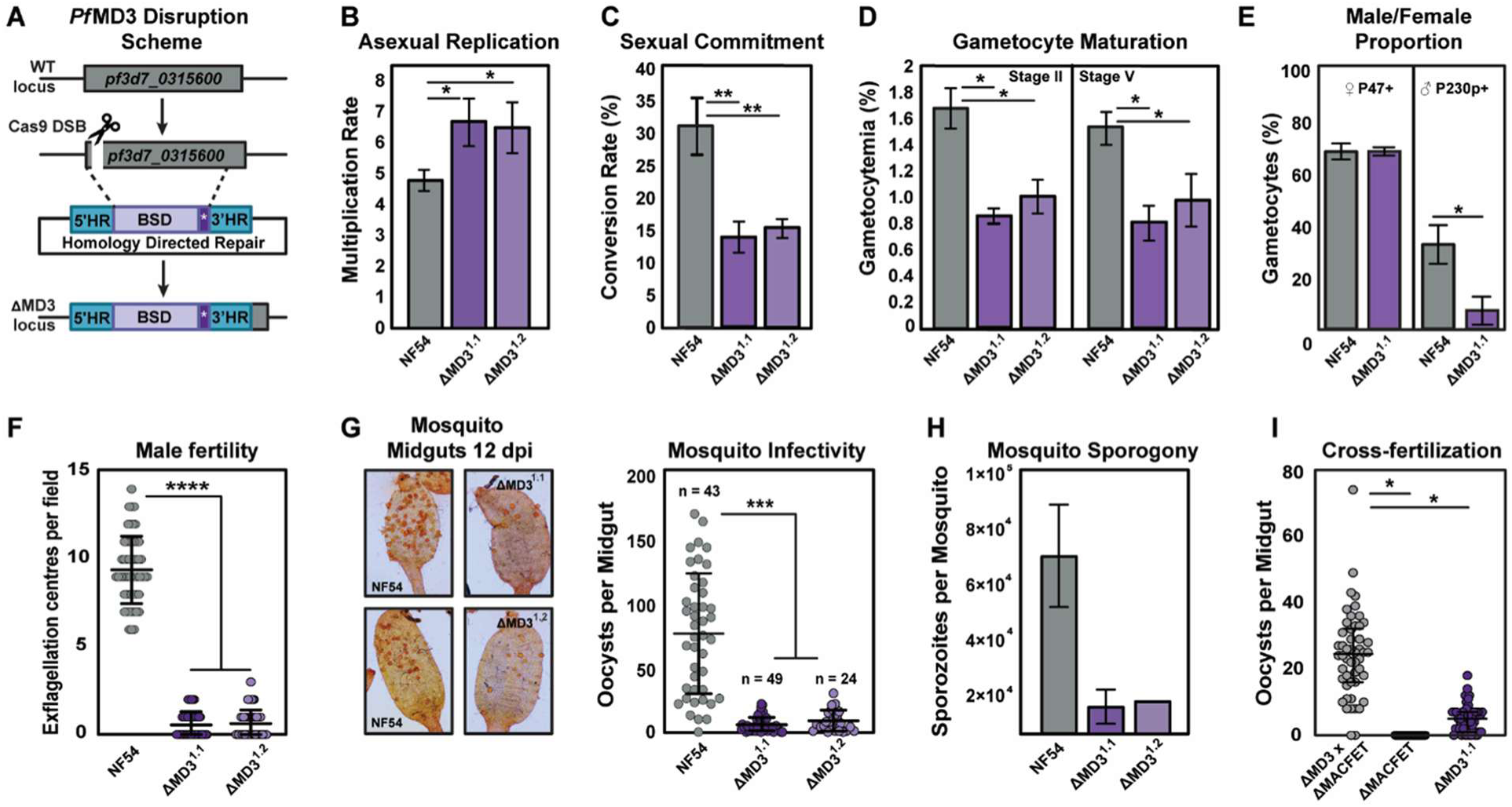
*Pf*MD3 plays an integral role in *P. falciparum* male fertility. **(A)** A CRISPR-Cas9 based approach was used to produce clonal genetic knockout lines of *Pf*MD3 by completely deleting the 1^st^ two exons and splice site of the genetic locus by using both a 5’ and 3’ homology region (HR) and swapping in a blasticidin (BSD) resistance cassette. Two clonal Δ*Pf*MD3 parasite lines were generated (1.1 and 1.2) and then assessed for **(B)** asexual blood stage replication rate, **(C)** percent gametocyte conversion rate, **(D)** Stage II and V gametocytemia quantified via preparation of Giemsa-stained thin-blood smears of *in vitro* cultures (n = 3, minimum 1000 cells per biological replicate) and **(E)** sex ratio determined by P47 and P230p staining (n = 2, at least 30 gametocytes counted per replicate). **(F)** The ability of Δ*Pf*MD3 to produce male gametes was investigated by inducing exflagellation of gametocytes (day 15) and counting the number of exflagellating centers per field (n = 3 for each clone). The infectivity of Δ*Pf*MD3 to *Anopheles stephensi* mosquitoes was determined by quantifying **(G)** midgut oocyst number on day 12 and **(H)** salivary gland sporozoites on day 14 after feeding. Data are shown for the NF54e parent (n = 2) and liPfMD31 1 (n = 2), with *p* = 0.1 and liPfMD31 2 (n = 1). **(I)** Δ*Pf*MD3 clone^1.1^ and the female-deficient ΔMACFET parasites^63^ were each fed to *Anopheles stephensi* mosquitoes at 0.3% gametocytemia and also mixed in a 1: 1 ratio (0 .3% gametocytemia) and fed to mosquitoes. Oocysts were quantified 10 days after infection (n = 2). Statistical analysis was performed with an unpaired, two-tailed t-test (B, C, D, and E, F, H, I) or one-way ANOVA (G), with significance indicated * = *p* < .05, ** = *p* < .01, *** = *p* < .001, **** = *p* < 0.0001 and error bars represent SEM.

To determine if the increased multiplication rate was due to a change in the proportion of asexual parasites compared to sexual gametocytes in the total population, we measured the ability of wild-type parasites versus the Δ*Pf*MD3^1.1/1.2^ clones to commit and progress through sexual development. Overall, the rate of commitment was reduced by 50% (Figure 2C) compared to the NF54e parental line. To further validate this observation, we also measured the multiplication rate for the NF54e and Δ*Pf*MD3^1.1^ lines grown in the presence of 2 mM choline chloride which is known to suppress sexual commitment^16^. Our results show that, in the presence of choline, there was no difference between the asexual multiplication rate in NF54e and Δ*Pf*MD3^1.1^ (Figure S2F). Therefore, we predict that the increase in multiplication we observe in Δ*Pf*MD3^1.1^ is due to a concomitant decrease in sexual commitment. To control for extraneous factors (i.e.: temperature, pH, genetic adaptation from clonal knockout selection) that could influence growth differences between the NF54e and Δ*Pf*MD3^1.1^ strains, we generated a clonal *Pf*MD3-3xHA-glmS parasite line (Figure S3A, B) for regulatable knockdown of PfMD3 expression in the presence of glucosamine^76^. Growth of *Pf*MD3-3xHA-glmS parasites in 1.25 mM glucosamine resulted in >90% knockdown of *Pf*MD3 expression after 72 h (Figure S3C) and phenocopied the growth phenotypes we observed in Δ*Pf*MD3 suggesting MD3 is important for enabling commitment to sexual development (Figure S3D).

Following sexual commitment, we observed a significant reduction in gametocytemia within the Δ*Pf*MD3 population (Figure 2D, day 10 post-induction, Stage IV-V gametocytes). We next determined if this decrease was sex-specific and found a notable decrease in the number of mature male gametocytes expressing P230p compared to wild-type, with no change in the proportion of P47-positive mature female gametocytes (Figure 2E). This suggests that the overall decrease in gametocytemia is due to a sex-specific impact of *Pf*MD3 in males. However, the mature Stage V Δ*Pf*MD3 parasites appeared morphologically similar to wild-type parasites (Figure S2G).

Because we observed a decrease in P230p positive male gametocytes in the Δ*Pf*MD3 line, we measured male fertility by quantifying exflagellation of mature Δ*Pf*MD3 male gametocytes. This fertility assay revealed that the average number of exflagellation centers were reduced ten-fold (Δ*Pf*MD3^1.1^ = 0.56 ± 0.10, Δ*Pf*MD3^1.2^ = 0.63 ± 0.10 mean ± SEM) compared to 9.40 ± 0.26 exflagellation centers per field in the NF54e parent (Figure 2F). Loss of male fertility was independently validated following *Pf*MD3-glmS knockdown as well (Figure S3E). To further define the male fertility defect, we measured the nuclear content of activated gametocytes using flow cytometry. This analysis revealed a significant reduction in male gamete DNA content (Figure S2H), suggesting a defect in the ability of the male gametocytes to properly replicate their genomes in preparation for exflagellation. Finally, to directly measure the transmission capacity of Δ*Pf*MD3 gametocytes, self-crossed Δ*Pf*MD3 or NF54e gametocytes were fed to *Anopheles stephensi* mosquitoes to determine the number of midgut oocysts at 12 days post feeding. As anticipated, Δ*Pf*MD3 clones produced approximately ten-fold fewer oocysts than the wildtype parental line (Figure 2G) with a corresponding reduction in day 14 salivary gland sporozoites (Figures 2H, S2I).

To further characterize the male defect in gametogenesis, we performed a genetic cross in *Anopheles stephensi* mosquitoes between Δ*Pf*MD3^1.1^ and the *P. falciparum* ΔMACFET^63^ strain that is deficient in the production of fertile females. As expected, self-crossed ΔMACFET produced no observable oocysts due to the lack of fertile females. However, when Δ*Pf*MD3 was crossed with ΔMACFET the resulting oocyst numbers were increased (24.4 ± 2.4 oocysts per midgut) compared to either Δ*Pf*MD3 or ΔMACFET self-crosses (Figure 2I). These results suggest that, despite the normal morphological development of the Δ*Pf*MD3 gametocytes, disruption of *Pf*MD3 reduces the production of reproductively competent male gametocytes.

### Bulk and single-cell transcriptomics reveal reduced male gametocyte marker expression in Δ*Pf*MD3

Since *Pf*MD3 is necessary to produce fertile male gametocytes, we used RNA-sequencing to measure changes in mRNA abundance between Δ*Pf*MD3^1.1^ and NF54e gametocytes at Stage II and V of development (5 and 11 days post gametocyte induction, respectively) (Table S1). Across these two timepoints, a total of 459 transcripts were significantly different (Figure 3A; *p <* .05, Log_2_ FC in 95^th^ percentile). Of these, 363 transcripts were decreased in one or both timepoints in Δ*Pf*MD3 compared to the wild-type, including 191 genes that have been previously associated with sex-differentiation^9^ (Figure 3A; Table S1). Two-thirds of these transcripts associated with sex-differentiation encode proteins involved in male gametocyte development (127/191, *p* < .0001, Fisher’s exact test) (Figure 3A; Table S1). When we compared the expression of male and female markers from two previous *P. falciparum* transcript datasets^9,15^ to our Δ*Pf*MD3 results, we found that male markers were significantly decreased in Stage V (*p < .*001) compared to female markers (Figure 3B). These changes in the transcriptome were most pronounced long after the peak of *Pf*MD3 expression at Stage II of gametocytogenesis.

**Figure 3.**
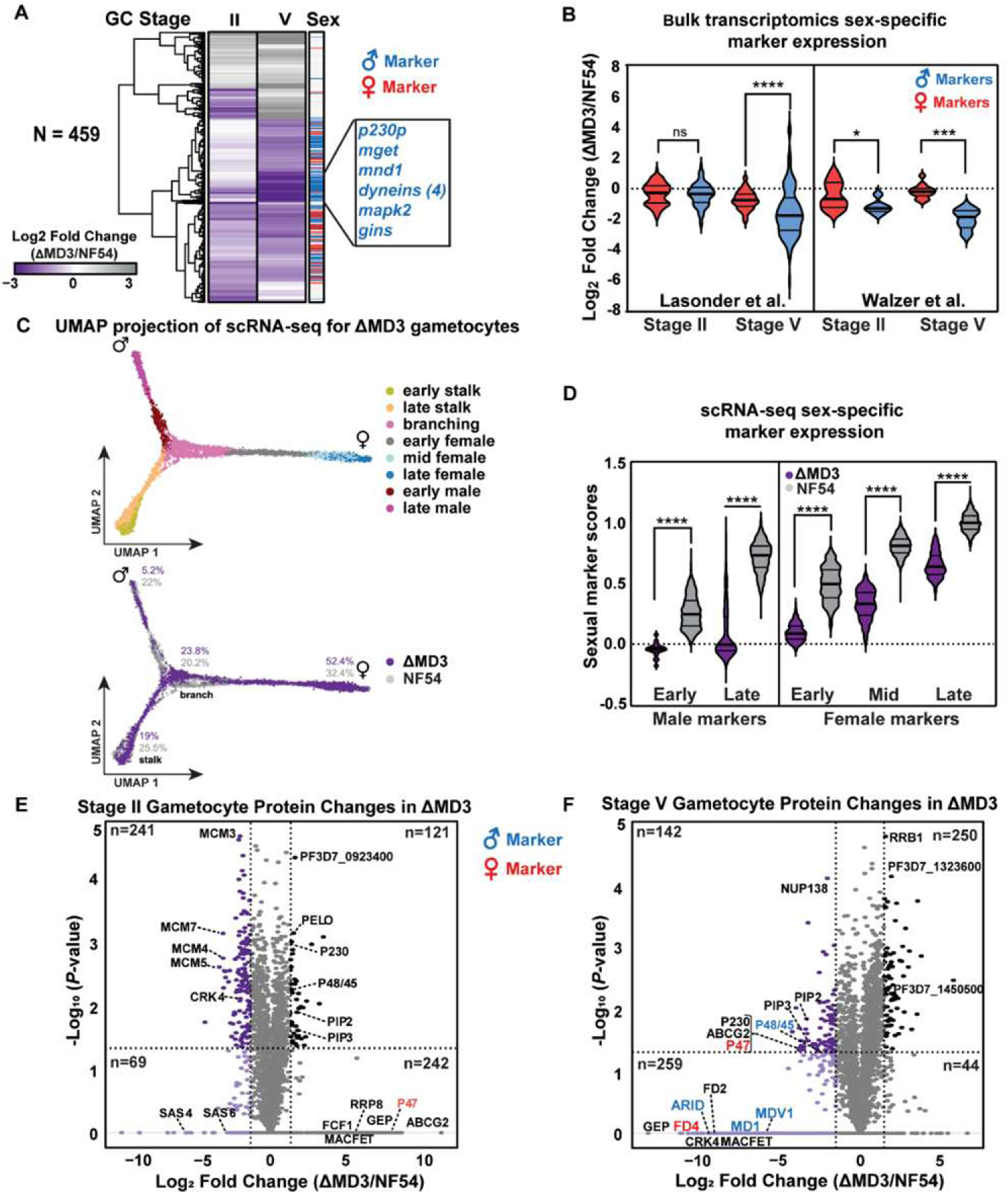
Altered transcript and protein abundance in Δ*Pf*MD3 is strongly associated with its male a development defect. (**A**) Differentially expressed transcripts were identified between *P.f.* strains Δ*Pf*MD3^1.1^ and the NF54e parent from bulk RNA-sequencing by using DEseq2 (*p* < .05) and Log_2_ Fold Change ± 1.5 for the average of the biological replicates (n = 2) with the associated data in Table S1. Transcripts displayed in the heatmap were clustered by Euclidean distance and known gametocyte gene markers were identified in each cluster (male markers in blue and female markers in red). (**B**) Data from bulk RNA-sequencing of separated male and female gametocytes (Lasonder)^9^ and single cell qPCR (Walzer)^15^ were used to select male and female-enriched transcripts. The Log_2_ Fold Change expression in Δ*Pf*MD3^1.1^/NF54 is represented in violin plots. Significant differences between male and female markers at each stage were determined with Welch’s t-test with significance indicated (* = *P <* .05, *** = *P <* .001, **** = *P <* .0001). (**C**) scRNA-seq UMAP trajectories of 3,106 subsetted cells from Δ*Pf*MD3^1.1^ that were integrated with the subsetted NF54 gametocyte Malaria Cell Atlas (7988 cells)^14^ (upper panel) with relative proportions of male, female and developing cells indicated on the UMAP graph (lower panel; Δ*Pf*MD3^1.1^ = purple and NF54 = grey). (**D**) scRNA-seq data of male and female markers from (B) are represented as the relative sexual score for both male and female clusters from Δ*Pf*MD3^1.1^ or NF54. Volcano plots show differentially expressed proteins in Stage II (**E**) or Stage V (**F**) gametocytes with thresholds indicated for the 95^th^ percentile (vertical lines) and *p <* .05 (horizontal lines).

To identify possible changes in specific subsets of sexual stage gametocyte populations (pre-differentiation, male, and female), we performed single cell RNA-seq (scRNA-seq) on Δ*Pf*MD3^1.1^ throughout gametocytogenesis from Days 0-12 (Figure 3C). We captured 7,026 high-confidence single cells spanning asexual and sexual developmental stages. To improve the resolution of post-commitment sexual development and enable direct comparison with wild-type parasites, we integrated our dataset with the *P.f.* strain NF54 scRNA-seq Malaria Cell Atlas from Dogga et al.^14^ (Figure S4D). Within the integrated datasets, the male cluster accounted for 5.3% of Δ*Pf*MD3 cells and 22.0% of NF54 cells, whereas the female cluster accounted for 75.7% and 52.5%, respectively.

We then evaluated cells classified into early gametocytogenesis as well as those assigned to a particular sex lineage across the integrated datasets using refined cluster annotations and pseudotime trajectory analysis (Figure S4E). When cells were classified by developmental stage (early/late) and sex using established markers^9^, Δ*Pf*MD3 cells consistently exhibited a lower sex-specific module score across all male and female gametocyte clusters compared to NF54, with late male gametocytes showing the greatest relative decrease (Figure 3D). This suggests that *Pf*MD3 influences sexual maturation, and its loss is most detrimental to male fertility.

Comparing bulk transcriptomics and scRNA-seq revealed that the largest overlap between the datasets was in the decreased male lineage transcripts (36%) compared to 9% in the female lineage in Δ*Pf*MD3. Specifically, 74% of decreased transcripts in the male lineage were male-specific genes^9,15^ (Table S7), suggesting a targeted disruption of the male-specific gene expression program beyond the overall decrease in male gametocytes. This suggests that while male gametocyte depletion drove bulk transcriptome changes, the few male gametocytes in Δ*Pf*MD3 captured by scRNA-seq showed intrinsic defects in male marker expression. To further characterize these male-specific transcriptional defects, we examined individual gene expression patterns and found significant decreases at multiple timepoints in both the bulk transcriptome and in the male lineage captured by scRNA-Seq, as exemplified by *mapk2: pf3d7_1113900* and *p230p: pf3d7_0208900* (Figure S4A; Table S7). In contrast, transcripts of female markers such as *p25* (*pf3d7_1031000*) and *pf47* (*pf3d7_1346800*) were not significantly affected by *Pf*MD3 deletion in any lineage or in bulk transcriptomics (Figure S4A). Additionally, essential non-sex-specific gametocyte markers such as *ap2-g* (*pf3d7_1222600*) and *pfs16* (*pf3d7_0406200*) showed no reduction in either transcriptome (Figure S4A). These results suggest that the morphologically defined male gametocytes in the Δ*Pf*MD3 population are transcriptionally aberrant and developmentally defective.

Following lineage bifurcation, a substantial set of genes remained differentially abundant within both male and female lineages. Transcripts with lower abundance in Δ*Pf*MD3 compared to NF54 were enriched for epigenetic and transcriptional regulators, including bromodomain protein 3 (*pf3d7_0110500*)^17^, ELM2 domain-containing protein (*pf3d7_0108500*)^18^, and multiple ApiAP2-domain containing proteins (*ap2-g2: pf3d7_1408200*^32^*; ap2-o2: pf3d7_0516800, ap2-g5: pf3d7_1139300*) (Figure S4F). Together, these changes suggest that *Pf*MD3 functions early in gametocytogenesis to establish a transcriptional program required to sustain chromatin accessibility and ApiAP2-mediated gene regulation. Loss of *Pf*MD3 thus disrupts expression of key transcriptional regulators, likely creating a cascade effect that impairs proper gametocyte maturation and differentiation.

### Proteomic analysis confirms diminished male gametocyte-specific protein levels in Δ*Pf*MD3

To complement our transcriptomic analyses, we measured total cellular protein abundance in Δ*Pf*MD3 compared to NF54 parasites at early-stage (Stage II, day 5) and late-stage (Stage V, day 11) gametocyte development (n = 2) using Data-Independent Acquisition Mass Spectrometry (DIA-MS) (Figures 3E, F; Table S2). DIA-MS proteomics revealed 763 proteins with differential abundance at Stage II (363 increased, 310 decreased; Figure 3E), with more pronounced changes at Stage V (401 decreased, 294 increased; Figure 3F). Importantly, 39 showed concordant transcript reductions in scRNA-seq analysis, the majority representing the transcriptional and epigenetic regulators identified above (Figure S4F), demonstrating that a handful of transcriptional changes translate to protein-level defects.

Among the reduced proteins were several that contribute to male exflagellation (Figure 3F; Table S2), including members of the 6-cysteine family that regulate gamete fertility (P230p, P230, P47, P48/45^77^) and several male-gametocyte enriched protein markers (ARID/MD4^61,78^, MD1^60,61^, MDV1^79^) (Figure 3F; Table S2). Interestingly, despite no overall reduction in the number of female parasites detected in culture, some proteins essential for female fertility were also decreased in late-stage Δ*Pf*MD3 gametocytes, including

MACFET^63^/FD1, FD2, and FD4^61^ (Figure 3F; Table S2). We also observed that 25 ribosome biogenesis-related proteins (GO:0042254) were increased in Δ*Pf*MD3 (Table S2). These include a ribosome assembly protein RRB1 (PF3D7_0816000) in Stage V gametocytes (Figure 3F) and pelota homolog PELO (PF3D7_0722100), which rescues stalled ribosomes^80^, as well as proteins involved in rRNA processing (RRP8: PF3D7_0925200, FCF1: PF3D7_0818400) in Stage II gametocytes (Figure 3E). The increase in ribosome biogenesis factors alongside decreased male and female fertility markers suggests that loss of *Pf*MD3 may disrupt regulation of gene expression during gametocyte development and that this process is essential to male gametocytes.

### *Pf*MD3 binds gametocyte mRNAs through GAA-repeat recognition

Since *Pf*MD3 contains a CCCH-type zinc finger domain and its disruption decreases male gametocyte-related gene expression, we investigated whether it binds to specific mRNA sequences^81–83^. To test this, we first measured whether *Pf*MD3 can bind directly to RNA using <u>C</u>ross <u>L</u>inking Immuno<u>p</u>recipitation followed by high-throughput <u>seq</u>uencing (CLIP-seq)^84,85^ at day 5 (Stage II) and 11 (Stage V) from *Pf*MD3-3xHA gametocytes (Figure 4A, Figure S5A). In total, we identified 207 mRNAs bound by *Pf*MD3, including 143 transcripts during Stage II and 86 transcripts at Stage V of gametocyte development, with 22 transcripts bound at both stages. Several of these mRNAs bound by *Pf*MD3 at both stages encode for proteins that are known markers of gametocytogenesis and have been demonstrated to play essential roles in gametogenesis (i.e., *p230: pf3d7_0209000, cdpk1: pf3d7_0217500*^86,87^) and transmission (*spm3: pf3d7_1327300*)^88^ (Figure 4A and Table S3). In addition, other regulators of male fertility (*md2: pf3d7_1233200, md4/arid: pf3d7_0603600*^61,78^) were bound at Stage II and Stage V respectively, while only a single regulator of female fertility (*fd3: pf3d7_1319600)* was bound in Stage V gametocytes, suggesting a bias of *Pf*MD3 to binding transcripts important for male development.

**Figure 4.**
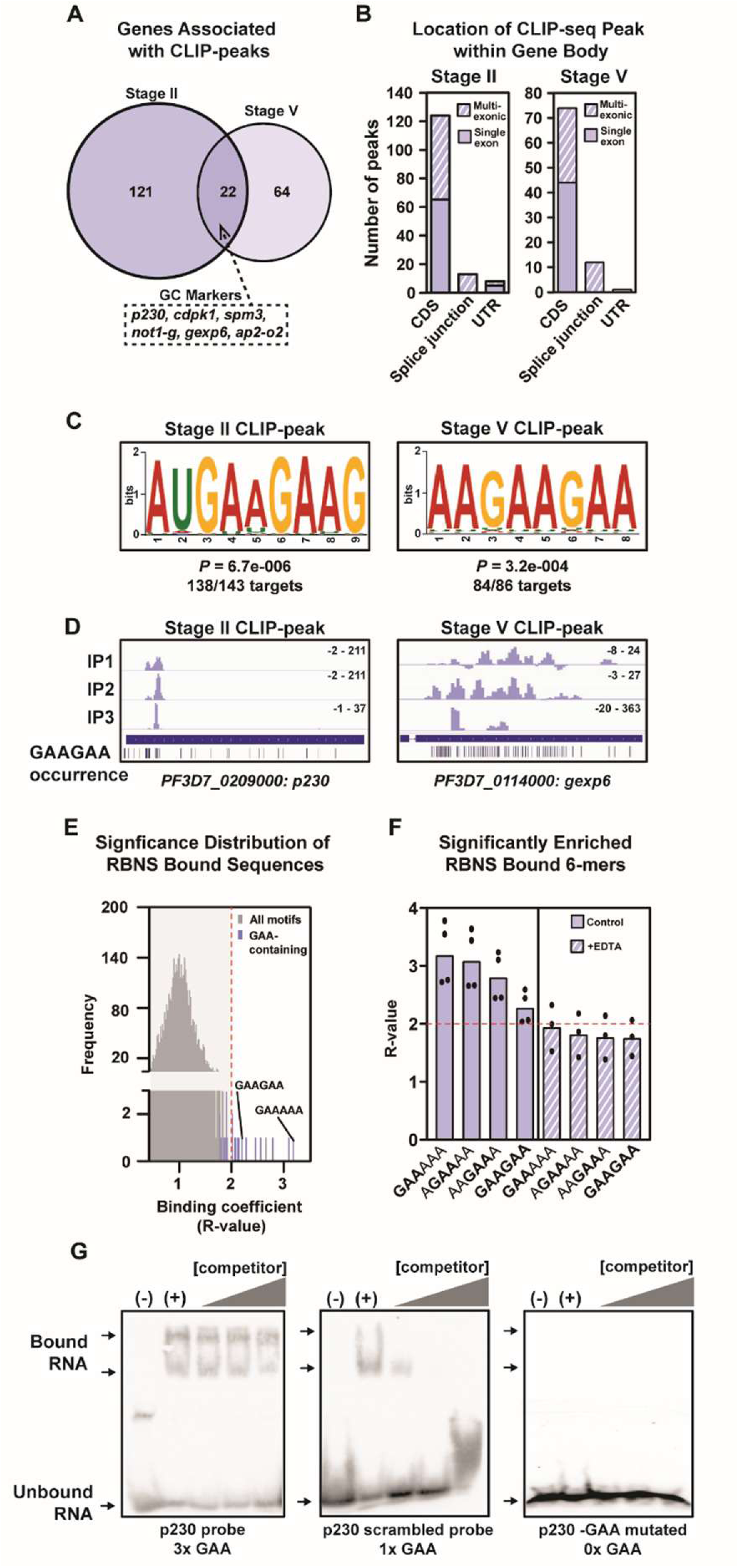
*Pf*MD3 interacts with RNA in gametocytes through binding a specific nucleotide sequence. CLIP-seq was performed using a *Pf*MD3-3xHA tagged line at Stage II and V of gametocyte maturation. Bound mRNA regions were identified by RNA sequencing combined with a bioinformatic pipeline employing Piranha^151^ (*p <* .01) and PureCLIP^150^ against input and IgG controls to call peaks. Peaks were retained if called in at least 2/4 biological replicates for each stage and visualized in IGV v2.1.3. (Associated data can be found in Table S3). (**A**) Venn diagram indicates overlap in bound transcripts called in each stage of development for a total of 207 bound transcripts. (**B**) Genomic features bound by *Pf*MD3 were binned into enriched regions spanning exons, across splice junctions and both exons and untranslated regions (UTRs) in genes containing single (solid) or multiple exons (hashed) and displayed as a bar graph. (**C**) The most enriched motifs within the bound transcripts were identified using STREME^92^ with the *P. falciparum* 3D7 reference genome v59.0^99^ as control sequences. (**D**) Examples of transcripts visualized in IGV v2.1.3. with the peak enrichment of reads in the IP following input subtraction and the location of each occurrence of the enriched motif for each stage. (**E**) RNA-bind-n-seq (RBNS) was used to identify all possible 6-mers bound by recombinant *Pf*MD3 CCCH-type zinc finger domain. Data are presented as enrichment (R-values) with the significance threshold indicated with dashed red lines. Several GAA-containing motifs are highlighted (purple). (**F**) Displayed are the R-values for the top motifs obtained by RBNS for *Pf*MD3 with 5 mM EDTA (hashed bars) or without (solid bars). (**G**) RNA EMSA measuring the binding of the recombinant CCCH zinc finger domain to either a probe from the P230 coding sequence containing 3x GAA, a scrambled version of the same sequence containing 1x GAA, or the P230 probe in which the three GAA instances were all mutated to CUU. Legend: (-) = no protein control, (+) = no competitor control, [competitor] = unlabeled RNA competitor probes were added in increasing concentration (10x, 100x, 1000x concentration compared to biotinylated control), whereas unlabeled scrambled probe was added to the middle EMSA and GAAGAA probe on the right.

To determine whether the RNAs bound by *Pf*MD3 encode proteins with functional roles relevant to gametocytogenesis, we evaluated all targets for GO-term^89^ enrichment (Table S3). At both stages tested, RNA/mRNA destabilization (GO:0050779 and GO:0061157) was the most significantly overrepresented function among bound transcripts (Table S3). Additionally, transcripts bound at Stage II in development were enriched for regulators of transcription (GO:0045893) and chromatin remodeling (GO:0006338) including 4 ApiAP2 proteins and transcripts bound in Stage V were enriched for protein kinases (GO:0016310) involved in post-translational regulation. These results suggest that the gene products bound by *Pf*MD3 include important regulators of gene expression that could aid in establishing the sexual developmental program.

The location of bound regions in mRNA transcripts can be indicative of RNA binding protein function^90^. Since our CLIP-seq demonstrated that *Pf*MD3 interacts with specific RNAs *in vivo*, we examined the distribution of *Pf*MD3 binding sites within individual mRNAs to determine whether it preferentially binds nascent RNAs. We found that the majority of *Pf*MD3 binding occurred within the exonic regions for both single and multi-exonic genes (85%, Figure 4B). *Pf*MD3 binding sites were rarely found across splice sites and within intronic sequences (Figure S5B). Intron splicing is therefore not likely the primary role of *Pf*MD3. However, the presence of some intronic sequences in bound transcripts indicates that binding may occur prior to or during mRNA processing for translation^62,91^.

To determine whether *Pf*MD3 interacts with mRNA in a sequence-specific manner, we performed motif enrichment analysis of the bound transcripts using STREME (Sensitive, Thorough, Rapid, Enriched Motif Elicitation; https://meme-suite.org/meme/doc/streme.html^92^). Strikingly, the vast majority of the bound transcripts contained a similar enriched motif, AWGAAGAA, at both Stage II (136/143 mRNA targets; *p =* 6.7 × 10^-6^) and Stage V (84/86 mRNA targets; *p =* 3.2 × 10^-4^) (Figure 4C), occurring at a frequency above the transcriptome-wide background (8415 occurrences in transcripts, www.Plasmodb.org). Closer examination revealed that this motif occurred in repeated clusters within 66 of the 207 enriched regions identified by CLIP-seq to be bound by *Pf*MD3 (Figure 4D, Figure S5C), suggesting that the AWGAAGAA sequence repeats may be preferred over single GAA trinucleotide occurrences.

### Dissecting the *Pf*MD3 ZnF-domain sequence-specific interaction

To directly test the sequence specificity of the RNA interaction with the *Pf*MD3 CCCH-type ZnF domain, we employed <u>R</u>NA-<u>B</u>ind-<u>N</u>-<u>S</u>eq (RBNS)^93^, an unbiased approach in which a recombinant protein domain is incubated with a random RNA library prior to pull-down and high-throughput sequencing of the bound RNA. This analysis confirmed the preferential binding of the *Pf*MD3 CCCH-ZnF domain to GAA-containing RNAs (Figure 4E). To ensure that sequence-specific binding depends on the structural integrity of the CCCH-type zinc finger domain, we disrupted the ZnF fold by adding EDTA, a zinc chelating agent, to the RBNS reaction. As expected, treatment with EDTA abolished sequence-specific binding (Figure 4F). Using the GAA triplet core sequence identified by RBNS to search within the *Pf*MD3 CLIP-seq peaks, revealed that 74% of peaks contained >1 instance of GAA, with over 90% having >10 instances (Figure S5D). Given that a single zinc finger domain typically only binds a triplet or quadruplet nucleotide sequence with high specificity^94^, these finding suggest that while individual GAA motifs are sufficient for binding, the extensive GAA repeats observed in target transcripts may facilitate binding by multiple *Pf*MD3 monomers or multimeric complexes.

To further test the interaction of the *Pf*MD3 CCCH-type ZnF domain, we used RNA <u>e</u>lectrophoretic <u>m</u>obility <u>s</u>hift assays (EMSA) with *p230*, a male gametocyte marker identified as a *Pf*MD3-3xHA target by CLIP-seq (Figure 4G). Using a biotin-labeled RNA probe designed from the *p230* coding sequence containing multiple GAA-repeats, we observed robust binding by the recombinant *Pf*MD3 CCCH-type ZnF domain that could not be outcompeted by a scrambled control probe at any concentration tested (Figure 4G). Interestingly, we also observed a supershift in the *p230* RNA probe, which may indicate multimerization of the CCCH-type zinc finger. We also performed EMSAs with a scrambled *p230* RNA probe retaining a single GAA trinucleotide and a mutant probe with all GAA instances replaced by CUU (Figure 4G). While the *p230* scrambled probe was weakly bound, the GAA-to-CUU mutant probe showed no detectable binding. We conclude that *Pf*MD3 binds a GAA-triplet motif via the CCCH-type ZnF domain and that these trinucleotide sequences often occur in repeated clusters within RNA transcripts bound by *Pf*MD3.

### *Pf*MD3 promotes translation of mRNAs required for male gamete production

CCCH-type ZnF RNA-binding proteins play a variety of roles in mRNA metabolism^83^ including translational repression, transcript maturation, decay^68^, and protein translation^95^. Perturbation of these processes would manifest as alterations to *Pf*MD3 bound mRNA transcripts, the levels of their protein products, or both. To determine the potential cellular role of *Pf*MD3, we examined mRNA and protein abundance changes between Δ*Pf*MD3 and wild-type NF54e for the 207 *Pf*MD3 mRNA targets identified from CLIP-seq (Figure 5A). If *Pf*MD3 primarily regulates mRNA stability or decay, we would expect to observe corresponding changes in transcript levels. Conversely, if *Pf*MD3 regulates translation, we would expect changes primarily at the protein level with minimal effects on transcript abundance.

**Figure 5.**
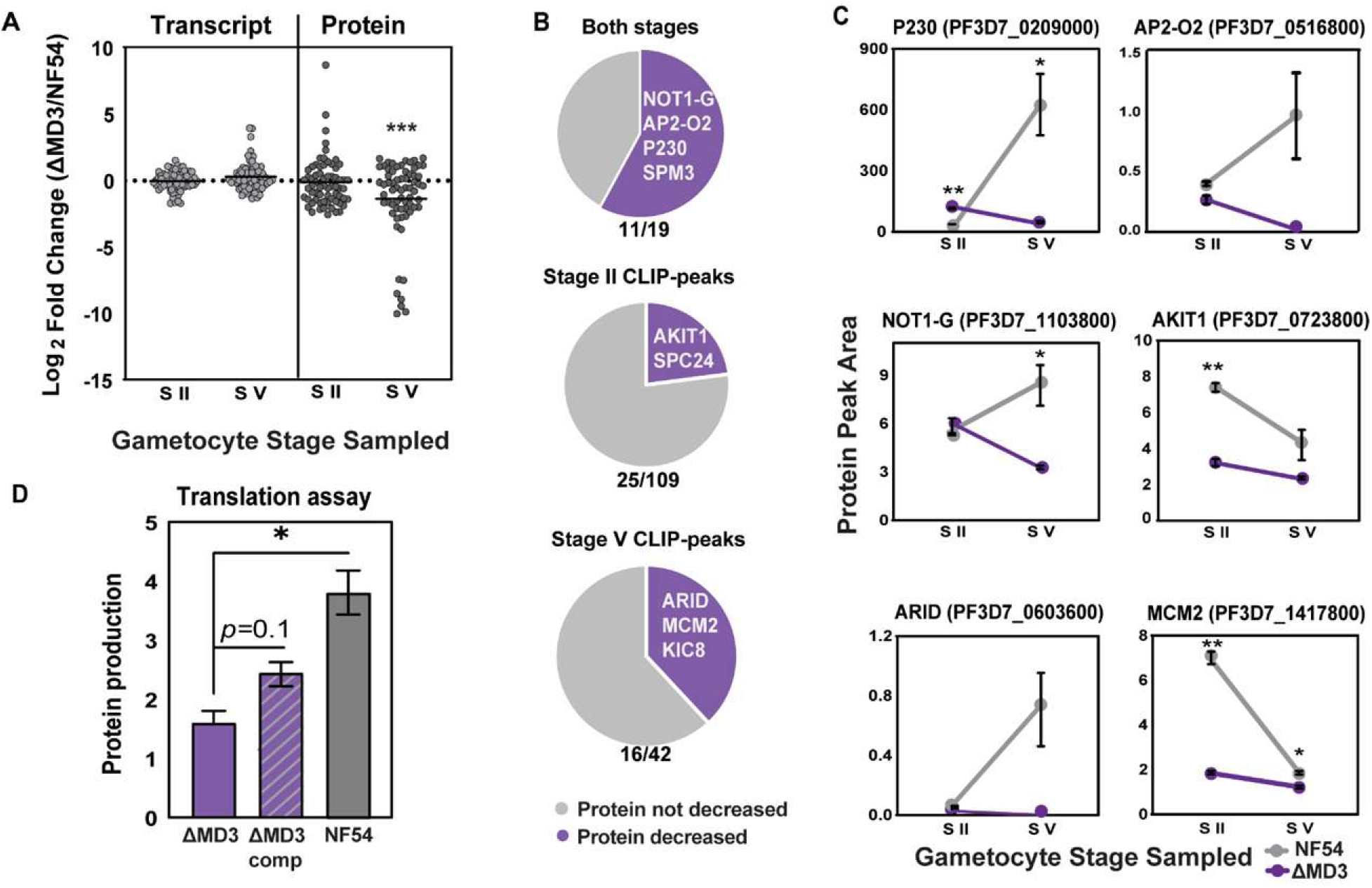
Protein abundance of *Pf*MD3 mRNA targets is altered in Δ*Pf*MD3. (**A**) Dot plot of the 66 differentially abundant *Pf*MD3 CLIP-seq mRNA targets and their relative Log_2_ Fold Change (Δ*Pf*MD3/NF54) from the transcript abundance and whole cell proteomics of Δ*Pf*MD3 at either Stage II or V of gametocyte development (Associated data in Table S4) with the black line representing the mean. A one-sample t-test was used to measure the difference of the mean from zero and significance indicated (*** = *p <* .001). (**B**) Pie charts of the proportion of CLIP-seq targets that are decreased (purple) in the Δ*Pf*MD3 proteome are shown in pie charts with specific examples indicated in slices. (**C**) Protein abundance of select genes of interest are plotted as line graphs (n = 2, mean ± S.E.M.) in NF54e and Δ*Pf*MD3 respectively and significant differences are indicated (Two-tailed t-test, * = *p <* .05, ** = *p <* .01). (**D**) The translation efficiency of either NF54 (grey), Δ*Pf*MD3 lysate (solid purple), or Δ*Pf*MD3 lysate with added recombinant *Pf*MD3-GST (*ΔPf*MD3 comp; hashed) was determined by an *in vitro* translation assay and protein production is plotted as a measure of protein concentration over time (fold change mg/ml from 0 h to 2 h) by BCA assay for n = 2 biological replicates (Two-tailed t-test, * = *p <* .05).

Surprisingly, most of the transcript targets of *Pf*MD3 show little to no change in the Δ*Pf*MD3 transcriptome at both Stage II and V (Figures 5A, S6A). In contrast, the protein products of these same targets were significantly decreased in Δ*Pf*MD3 compared to wild-type parasites (*p<.*001, Figure 5A) with 24 proteins reduced at Stage II and 27 proteins reduced at Stage V (Figure S6A; Table S4). This discordance between transcript and protein levels strongly suggests that *Pf*MD3 functions primarily at the translational level rather than regulating mRNA stability.

The greatest decrease in protein levels occurred for RNA targets bound by *Pf*MD3 at both stages Stage II and Stage V of gametocyte development, with over 50% of total targets significantly decreased (Figure 5B). These dual-stage targets include key regulators of male gamete development: P230^77^, SPM3^88^, NOT1-G^96^ and the transcription factor AP2-O2, which is necessary for male gametocytes gene expression^21^ (Figure 5C). Additionally, 23% of Stage II CLIP-seq targets showed decreased protein abundance in Δ*Pf*MD3 parasites, including kinetochore components (ex: AKIT1: PF3D7_0723800), while 38% of Stage V targets were decreased, including DNA replication factors (ex: MCM2: PF3D7_1417800) (Figure 5B, C, S6A; Table S2). Critically, among the targets of *Pf*MD3, several proteins with validated roles in male gamete formation were undetectable in Δ*Pf*MD3 stage V gametocytes: ARID/MD4^78^, AP2-O2^21^, KIC8 (PF3D7_1014900), and SNF2 ATPase (PF3D7_0216000)^97^; Figure 5C; Table S2). This progressive protein depletion during gametocyte maturation reflects the cumulative impact of *Pf*MD3 loss on the translation of essential developmental regulators that are necessary for male fertility.

A previous study suggested that *Pf*MD3 functions as a translational repressor during gametocytogenesis^73^. To test this hypothesis, we examined whether *Pf*MD3 target mRNAs are translationally repressed in wild-type parasites by determining the fraction of CLIP-seq targets detected as proteins in wild-type NF54e Stage V gametocytes. Contrary to the repression model, DIA-MS detected protein products for 170/207 (82%) of the *Pf*MD3 mRNA targets (Table S2) demonstrating that these transcripts are actively translated in the presence of *Pf*MD3. To test whether *Pf*MD3 enhances translation, we performed *ex vivo* translation assays using Δ*Pf*MD3^1.1^ or NF54 gametocyte lysate (Figure 5D). Compared to wild-type NF54 lysate we measured a significant reduction in protein synthesis in Δ*Pf*MD3^1.1^ lysate (Figure 5D). Supplementation of Δ*Pf*MD3^1.1^ lysate with recombinant *Pf*MD3-GST protein restored protein production to levels approaching wild-type, demonstrating that the translational defect can be attributed the loss of *Pf*MD3 (Figure 5D). Taken together, these data establish that mRNA transcripts bound by *Pf*MD3 are translated into proteins and are not degraded or stabilized. The expression of *Pf*MD3 in gametocytes thus allows for the sufficient production of proteins involved in male gamete production and suggests that the accumulation of these proteins during gametocyte development is crucial prior to the rapid and energy-intensive process of male gamete production^98^.

### *Pf*MD3 protein interactors support its role in translation of mRNA targets

To further elucidate the molecular role of *Pf*MD3 in sexual development, we identified protein interactors of *Pf*MD3 at the peak of its expression in Stage II-III (day 5) gametocytes using immunoprecipitation followed by data independent acquisition (DIA) mass spectrometry (IP-MS) from the *Pf*MD3-3xHA tagged parasite line. Across three biological replicates, we detected 102 proteins significantly enriched in the *Pf*MD3-3xHA immunoprecipitants compared to nonspecific IgG antibody controls (Table S5). Of these enriched proteins, 37 were annotated with putative roles in mRNA metabolism, binding, or processing and 23 were predicted to localize to organelles involved in mRNA translation including pre-ribosome, nucleolus, and cytoplasmic ribonucleoprotein granules^99^ (Figure 6A; Table S5). Two interactors, both zinc finger domain containing proteins (ZNF4: PF3D7_1134600 and RNF1: PF3D7_0314700), were previously identified and verified via reciprocal immunoprecipitation^73^, providing independent validation of our IP-MS approach (Figure 6B).

**Figure 6.**
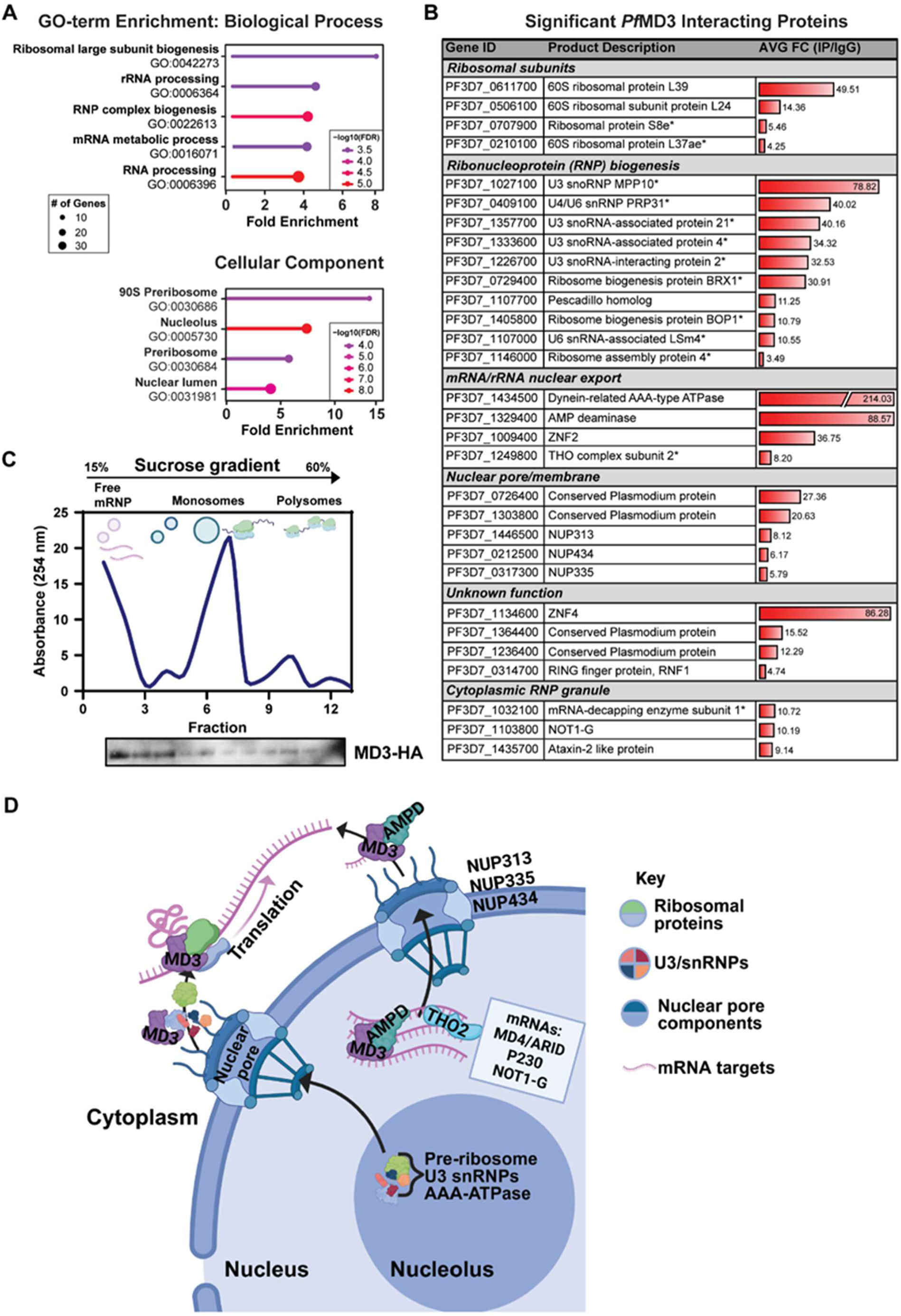
Protein interactors of *Pf*MD3-3xHA suggest a role in translational processing. (**A**) GO-term enrichment (FDR *<*0.05) of biological processes and cellular components for the 102 proteins found associated with *Pf*MD3-3xHA from IP-MS (n = 3) (associated data in Table S5). (**B**) PfMD3-interacting proteins of interest (* = putative annotation) with average enrichment fold change (HA/IgG) represented as data bars with numerical units noted. (**C**) Sucrose gradient fractionation of ribosome components with relative positions of free mRNPs, monosomes and polysomes indicated. The presence of *Pf*MD3-3xHA in each fraction was measured by western blot analysis. Shown is a representative western blot of n = 2 replicates, where replicate 2 is included in Figure S6E. (**D**) Collectively, this data supports a model where *Pf*MD3 functions at multiple levels of gene expression regulation. *Pf*MD3 may contribute to general translational capacity through interactions with U3 snoRNP complexes and ribosome biogenesis factors in the nucleolus, facilitating rRNA maturation or ribosome assembly. Additionally, *Pf*MD3 exhibits mRNA-specific functions: it binds directly to target mRNAs containing GAA-repeat motifs and associates with mRNA export machinery, including AMP deaminase, THO2, and nucleoporins (NUP313, NUP335, NUP434), suggesting it may escort specific transcripts from the nucleus to the cytoplasm. In the cytoplasm, *Pf*MD3 associates with free mRNPs and monosomes, ultimately promoting translation of its target mRNAs. It is unclear whether individual *Pf*MD3 molecules accompany specific mRNAs throughout their journey from nucleus to ribosome, or whether distinct pools of *Pf*MD3 function independently in ribosome biogenesis and mRNA-specific regulation (created using BioRender).

We also detected 21 proteins involved in nucleolar ribonucleoprotein (RNP) biogenesis (GO:0022613; Figure 6A, B; Table S5), suggesting that *Pf*MD3 participates in assembling translational machinery for protein production. To determine whether *Pf*MD3 associates with actively translating ribosomes, we profiled ribosomal fractions and found *Pf*MD3 primarily associates with free messenger ribonucleoproteins (mRNPs) and monosomes rather than the polysomes, which are typically associated with active translation (Figure 6C, S6E). This distribution pattern suggests that *Pf*MD3 engages with mRNAs prior to or during the initiation of translation by direct association with the translational machinery.

*Pf*MD3 also interacts with four members of the U3 small nucleolar RNP (U3 snoRNP) complex: MPP10, U3 snoRNA-associated proteins 21, 4, and 2. Additionally, *Pf*MD3 co-immunoprecipitated with two proteins of unknown function (PF3D7_1236400 and PF3D7_1364400) that were previously shown to interact with U3 snoRNP complex members in a yeast two-hybrid assay^100^ (Figure 6B). While U3 snoRNPs are well-characterized for their roles in rRNA maturation^101^, they also shuttle between the cytoplasm and nucleus of eukaryotic cells^102^ mirroring the dual localization observed for *Pf*MD3 (Figure 1C). This parallel localization pattern and evidence of interaction support a functional relationship between *Pf*MD3 and the nucleolar U3 snoRNP complex.

The two most highly enriched *Pf*MD3 interactors were a dynein-related AAA-type ATPase (PF3D7_1434500) and AMP deaminase (PF3D7_1329400) (Figure 6B, D). The dynein-related AAA-type ATPase is an ortholog of Midasin (41% sequence identity, 65% positive homology to human MDN1; https://blast.ncbi.nlm.nih.gov/Blast.cgi)^103^ which functions as a nuclear chaperone for 60S pre-ribosomal subunits in higher eukaryotes. AMP deaminase, conversely, has been implicated in gene gating and mRNA export in other systems^99,104^. We validated these interactions *in vitro* by protein pull-down of *Pf*MD3-3xHA from parasite lysate with recombinant AMP deaminase (Figure S6D) and MPP10 (Figure S6C), supporting the relevance of these interactions and *Pf*MD3’s proposed role in mRNA export.

Beyond ribosome biogenesis factors, our IP-MS identified additional interactors implicated in mRNA export. These include ZNF2 (PF3D7_1009400), an orthologue of a protein involved in nuclear export of mRNA in *Toxoplasma gondii*^105^; THO2 (PF3D7_1249800), a component of the transcript export complex^106^ (Figure 6B); and three nucleoporins—NUP313 (PF3D7_1446500), NUP335 (PF3D7_0317300), and NUP434 (PF3D7_0212500)—that form channels in the nuclear envelope (Figure 6B, D). The presence of these mRNA export machinery components among *Pf*MD3 interactors suggests that *Pf*MD3 could accompany its target mRNAs from the nucleus to the cytoplasm. Collectively, these data reveal that *Pf*MD3 is a multi-functional RNA-binding protein (Figure 6D).

## Discussion

*Plasmodium falciparum* undergoes an extended ∼14-day sexual maturation process, differentiating through five developmental stages, producing male and female gametocytes. Beyond Stage III of development, protein and RNA synthesis largely cease, and gametocytes enter a developmentally arrested state while circulating in the blood stream^107–110^, remaining poised for up to 21 days until female mosquito injection triggers activation^111^. Preparation for this poised phase likely relies on post-transcriptional regulation^6,31,32,112^. To fulfill these needs, hundreds of predicted RNA-binding proteins (RBPs) encoded in the *Plasmodium* genome are thought to play critical regulatory roles as transcription decreases during gametocyte maturation^33,35–37^.

The best-characterized Plasmodium RBPs—DOZI, CITH, and the Puf family proteins—function as translational repressors in female gametocytes, storing mRNAs for activation upon delivery to the mosquito^42–45^. In contrast, post-transcriptional regulation in male gametocytes likely differs substantially, as the rapid process of micro gametogenesis (∼15 minutes) necessitates that proteins required for gamete formation be translated and accumulated during the preceding maturation period rather than during exflagellation itself. Recently, three putative RBPs (MD1, MD3, and MD5) have been shown to function in maintaining male gamete fertility^60,61,73^. In this study, we demonstrate that *Pf*MD3 (PF3D7_0315600) is a CCCH-type ZnF RNA-binding protein essential to male gamete fertility through translational activation of proteins required for male gametocyte development.

### PfMD3 functions early in the male gametocyte developmental hierarchy

*Pf*MD3 is expressed throughout the intraerythrocytic life cycle with maximal abundance in Stage II gametocytes, positioning it as an early regulator of sexual development. Consistent with previously published phenotyping^73^, genetic disruption of *Pf*MD3 (Δ*Pf*MD3) reduced the proportion of functional male gametocytes and severely impaired exflagellation. By crossing Δ*Pf*MD3 with a fertile female deficient line, Δ*Pf*MACFET^63^ we could rescue fertility and confirm the male-specific function of *Pf*MD3. Using integrated transcriptomics and proteomics revealed that loss of *Pf*MD3 disrupts male lineage development, resulting in a drastic reduction of all later vector stages of development beyond oogony.

While *Pf*MD3 disruption primarily affects male development, we observed modest decreases in some female-enriched proteins (FD4, G377) in late-stage gametocytes. These likely represent secondary effects of early regulatory disruption rather than direct female-specific functions, as female gametocyte numbers remain unchanged and female gametes retain fertility. The translational repressor GD1, which binds both the md3 transcript and female-specific transcripts such as pfg27/25^72^, may normally repress md3 translation in female gametocytes where translational repression is the predominant regulatory mechanism. Loss of MD3 could subtly affect the balance of these interconnected regulatory networks during the critical Stage II period when sex-specific programs are established, creating minor effects on female development without compromising female gametocyte maturation or fertility.

### Sequence-specific RNA binding and translational activation

Using complementary *in vivo* (CLIP-seq) and *in vitro* (RBNS, EMSA) approaches, we demonstrated that *Pf*MD3 specifically binds GAA-trinucleotide repeats in RNA. The ubiquitous distribution of GAA-motifs throughout the *P. falciparum* genome^99^ suggests that developmental stage-specific expression of *Pf*MD3, combined with active transcription of its mRNA targets, confers specificity for male-specific transcripts.

Critically, loss of *Pf*MD3 resulted in a substantial reduction of target protein products with negligible change in the abundance of the mRNAs, revealing a translational role for *Pf*MD3 rather than transcriptional. This conclusion is further supported by: (1) IP-proteomics identifying *Pf*MD3 interactions with nuclear export machinery, ribosome biogenesis factors, and ribosomal subunits; (2) *Pf*MD3 association with free mRNAs and monosomes rather than polysomes; (3) detection of 82% of *Pf*MD3 mRNA targets as proteins in wild-type parasites, with decreased protein production in both the whole cell proteomics and *ex vivo* translation assays upon *Pf*MD3 loss. Our proposed role for *Pf*MD3 in interacting with translational machinery and other zinc finger proteins (ZnF4, RNF1) is consistent with published data for *Pf*MD3^73^. However, our findings do not support a role for *Pf*MD3 as a translational repressor as previously reported^73^. Rather, our multi-omics analyses and complementary *in vitro* methods demonstrate *that Pf*MD3 targets are actively translated and that loss of *Pf*MD3 decreases protein production.

Our data suggests that *Pf*MD3 functions at multiple levels: contributing to the general translational capacity through interactions with U3 snoRNPs and ribosome biogenesis factors, while promoting mRNA specific translation through direct GAA-repeat binding and facilitation of nuclear export and ribosomal engagement. This multifunctionality has precedent in mammalian systems, as human ZNF277 binds similar GAAGA motifs and regulates multiple aspects of mRNA metabolism through interactions with diverse protein complexes^113^. Whether individual *Pf*MD3 proteins accompany specific mRNAs from the nucleus to ribosomes, or whether distinct pools of *Pf*MD3 function independently in ribosome biogenesis and mRNA-specific regulation remains to be determined. This dual-function model positions *Pf*MD3 as a coordinator of both general translational capacity and mRNA-specific post-transcriptional regulation, ensuring proper expression of proteins essential for male gametocyte development.

### Hierarchical model of male gametocyte regulators

Our results position *Pf*MD3 within an emerging regulatory hierarchy that unfolds across multiple stages of sexual development. During the early sexual Stages I-II, *Pf*MD1 acts as the primary male-determining factor, necessary and sufficient for male fate commitment. *Pf*MD3 expression peaks at Stage II functioning downstream of or parallel to *Pf*MD1 and is required for male development^60^. From Stage II-V, *Pf*MD3 promotes the translation of key regulators including AP2-O2 (a transcription factor essential for male-specific gene expression), ARID/MD4, and NOT1-G. The decreased AP2-O2 protein level demonstrates translational control, potentially creating a feed-forward cascade where *Pf*MD3-mediated translation of transcription factors amplifies male-specific programs. During this same Stage II-V period, *Pf*MD5 may function to ensure proper splicing of male fertility transcripts, coordinating with *Pf*MD3 to ensure that correctly processed mRNAs are efficiently translated. Other MD proteins may function in later maturation or in gamete-specific differentiation^40,61,78^.

This model also captures the progressive protein depletion observed in ΔPfMD3 parasites, where 23% of Stage II targets and 38% of Stage V targets are affected, reflecting cumulative translational defects over developmental time. The complete loss of late-stage proteins such as AP2-O2 and ARID/MD4 despite mRNA presence demonstrates that *Pf*MD3-mediated translation is essential for accumulating these regulators to functional levels. The resulting developmental arrest and failed genome replication reflect insufficient accumulation of DNA replication machinery during the preparatory maturation phase, when these critical components should be building up in anticipation of later developmental events.

In conclusion, we establish *Pf*MD3 as a sequence-specific translational activator functioning in early gametocyte development and production of fertile male gametocytes. While our multi-omics approach demonstrates that *Pf*MD3 promotes translation of male gametocyte proteins; however, the precise mechanism of translational enhancement remains unclear. Our data suggest dual functions—contributing to general translational capacity through ribosome biogenesis while promoting mRNA-specific translation—but these have not been experimentally separated. Despite these questions, our work establishes that post-transcriptional regulation in male gametocytes fundamentally differs from the translational repression model in female gametocytes, reflecting distinct biological demands: females store mRNAs for rapid activation, while males accumulate proteins during extended maturation to support rapid exflagellation. A deeper understanding of these sex-specific strategies may reveal new transmission-blocking intervention targets.

## Acknowledgements

We thank Bryan Zavala (FDA, CBER), Dr. Tori Bonnell, and Dr. TJ Russell at the Pennsylvania State University for their contributions to this study. We would also like to acknowledge Dr. Kim Williamson (Uniformed Services University of the Health Sciences) for kindly providing us with anti-P230p antibody and Dr. Alvaro Molina-Cruz (Laboratory of Malaria and Vector Research, NIAID) for kindly providing us with anti-P47 antibodies. We also thank Drs. Stefan Kappe (Seattle Children’s Research Institute) and Sudhir Kumar (Iowa State University College of Veterinary Medicine) for providing their *P. falciparum* Δ*macfet* parasite line for performing genetic crosses. We would also like to thank Dr. Marcus Lee for providing the CRISPR-Cas9 plasmid backbone used for genetic modification of *Pf*MD3. We would like to acknowledge Drs. Wells Wu, Chao-Kai Chou and Rongfong Shen at U.S. FDA Center for Biologics and Evaluation Research, Facility for Biotechnology Resources and Drs. Haiyan Zheng and Caifeng Zhao at the Rutgers School of Medicine Mass Spectrometry Core Facility for processing samples related to this project. We acknowledge the critical resource the VEuPathDB project provides for accessing and analyzing *Plasmodium* data. Lastly, we would like to thank Dr. Godfree Mlambo, Chris Kizito, and the rest of the Insectary and Parasitology Core Facilities team at the Johns Hopkins Malaria Research Institute for their outstanding work. This work was supported by the Intramural Research Program of the Center for Biologics Evaluation and Research, Food and Drug Administration. The following funding resources were used: National Institutes of Health R01AI132359 (PS); Bloomberg Philanthropies (PS, AT, GX, and AA) and R01-AI125565 to ML and RVB, National Institute of Allergy and Infectious Diseases (NIAID) R01AI148489 to KS.

## Resource Availability

### Lead contact

Further information and requests for resources and reagents should be directed to and will be fulfilled by the lead author, Heather Painter.

### Materials availability

Parasite lines generated in this study (Δ*Pf*MD3 clones, *Pf*MD3-3xHA, *Pf*MD3-3xHA-glmS as well as the associated wild-type strains) are available upon request to the lead contact.

### Data and code availability

For genomic sequencing and single-cell RNA-sequencing results, raw reads were submitted to <u>SRA:PRJNA1111533</u>. For experiments that generated RNA, files were submitted to GEO: RNA-seq experiments, raw fastq.gz and processed Kallisto quantification files, accession number: <u>GSE267361</u> and for CLIP-seq experiment raw fastq.gz and processed bigwig files, accession number: <u>GSE267362</u>. Data from proteomics and IP-MS experiments were submitted to the MassIVE repository (https://massive.ucsd.edu/) with the identifier MSV000094788 and ProteomeXchange (http://www.proteomexchange.org) with the identifier PXD052415. Original western blot images for are included in supplementary figures and/or source data file and microscopy data used in this publication are available by request to the lead author.

## Methods

### Phylogenetic analysis and protein sequence alignments

Protein sequences of *Plasmodium* and *Hepatocystis* MD3 orthologs and syntenic genes were obtained from www.Plasmodb.org^99^. Protein alignments were performed using MUSCLE (MUltiple Sequence Comparison by Log- Expectation) available at https://www.ebi.ac.uk/Tools/msa/muscle/ and a distance corrected maximum likelihood phylogenetic tree of the protein alignments was constructed using interactive tree of life (iTOL) https://itol.embl.de/tree. Protein sequence analyses were conducted using Interproscan available at https://www.ebi.ac.uk/interpro/search/sequence/, NCBI conserved domain search with default settings available at https://www.ncbi.nlm.nih.gov/Structure/cdd/wrpsb.cgi. Nucleolar and nuclear localization signals in the *Pf*MD3 protein sequence were identified using NoD (<u>N</u>ucleolar localization sequence <u>D</u>etector)^74^ and NLStradamus^75^, respectively.

### Parasite line generation

To create a genetic knockout and tagged versions of *pf3d7_0315600* (*Pfmd3)* in *Plasmodium falciparum* NF54 Patient Line E (NF54e; obtained from BEI Resources (MRA-1000)), complimentary nucleotides encoding guide RNA on the 5’ end (for the knockout) and 3’ end (for the 3xHA tag) were selected using the CRISPR design function on Benchling (https://www.benchling.com) and evaluated for off-targets using Casoffinder^114^. The guide RNAs were inserted into the single plasmid for *P. falciparum* CRISPR/Cas9 editing, pDC2-U6A-hDHFR (kind gift of Marcus Lee of the Wellcome Sanger Institute, UK) using the BbsI site (reviewed in^115^). Homology regions representing the last exon and region upstream of the coding region were used to replace the first two exons and splice site of the *pf3d7_0315600* gene with a Blasticidin S Deaminase gene and stop codon. For the plasmid construct to introduce a 3xHA-tag, a homology region corresponding to the 3’ end of the gene, a 3xHA-tag followed by the stop codon and 3’UTR were included. These homology regions were inserted into pDC2-U6A-hDHFR using the EcoRI and AatII sites resulting in the final plasmid *p*DC2-U6A-hDHFR-MD3HA.

To create a glmS-knockdown version of *Pf*MD3 in the same NF54e parent line, the same homology region corresponding to the 3’ end of *pf3d7_0315600* used for the 3xHA-tagged line without the 3’UTR or stop codon was cloned into the pSLI-HA-glmS vector using XmaI and NotI restriction sites (kindly provided by Dr. Ron Dzikowski, the Hebrew University of Jerusalem).

Using the above-described plasmids, a tagged, glmS-knockdown or knockout transgenic line was created by electro-transformation into NF54e as previously described^116^. In brief, 100 µg of ethanol-precipitated plasmid in Cytomix (120 mM KCl, 0.2 mM CaCl_2_, 2 mM EGTA, 10 mM MgCl_2_, 25 mM HEPES, 10 mM K_2_HPO4/KH_2_PO4; pH 7.6) was transfected into NF54e ring-stage *Plasmodium falciparum* parasites using a Bio-Rad GenePulser set at 0.31 kV, 960 µF. The cultures were maintained for 48 h post-transfection at 4% hematocrit before adding media containing 2.5 nM WR99210 (a kind gift from Jacobus Pharmaceutical, Princeton, NJ, USA). For the glmS-knockdown transfection, drug selection was changed to neomycin for two weeks (G418; Sigma-Aldrich) at 400 µg/mL following appearance of WR99210 resistant parasites in culture. Clonal parasite lines were generated via limited dilution cloning, where parasites were seeded out to 0.3 parasites/well in a 96-well plate, monitored for growth over 14 days and subsequently genotyped (see Parasite genotyping described below), after which time parasites were maintained in drug-free RPMI medium.

### Parasite genotyping

Once stably growing parasites were observed in culture, parasite genomic DNA was extracted using the Qiagen DNeasy Blood and Tissue kit and genotyping PCR performed to verify the presence of modified parasites (Figures S1B, S2A, S2F), primer sequences provided in Table S6). Positive clones were also verified via whole genome sequencing performed as previously described^117^. Sequence output files were uploaded to Galaxy where the sequence reads were mapped to PF3D7 genome version 59.0 using BWA-MEM2 v2.2.1^118^. For the *Pf*MD3-HA-glmS line, whole genome sequencing was performed as per manufacturer’s guidelines using a MinION MK1C, where BAM files were basecalled in duplex with Dorado v0.5.1 using the super accurate model. BAM files were filtered to remove reads less than 200 bp and Q scores <20 prior to alignment to the *P. falciparum* 3D7 reference genome version 66.0. Sequence variants were identified using Freebayes (v1.3.8)^119^ prior to using SNPeff (version 5.2)^120^ to annotate and determine the statistical significance of changes from the reference genome.

### Parasite culturing and growth phenotyping

NF54e parasites and parasite strains generated for this study were cultured *in vitro* at 4% hematocrit in O+ human erythrocytes at 3–5% parasitemia as previously described^121^ in RPMI 1640 medium supplemented with 25 mM HEPES, 0.2% D-glucose, 200 μM hypoxanthine, 0.2% sodium bicarbonate, 24 μg/ml gentamicin (Biobasic, GB0217-5) with 0.5% AlbuMAX® I and incubated at 37 °C under hypoxic conditions (90% N_2_, 5% O_2_, and 5% CO_2_). For growth phenotyping complementation experiments using the *Pf*MD3-3xHA-glmS knockdown line, parasites were grown identically to the knockout and wild-type lines, with the addition of 1.25 mM glucosamine over 72 h prior to measuring relevant phenotypic endpoints.

### *Pf*MD3 knockout growth phenotyping

Following generation of clonal parasite knockout lines (Δ*Pf*MD3^1.1^ and Δ*Pf*MD3^1.2^), these parasites were studied for their asexual and sexual development *in vitro*.

#### Asexual development phenotyping

The parent NF54e parasites and Δ*Pf*MD3^1.1^ and Δ*Pf*MD3^1.2^ were seeded into 24-well plates (1 ml per well, 3 biological replicates) at 1% ring-stage parasitemia, 3% HC following 2 rounds of 5% D-sorbitol synchronizations, 6-8 h apart for two complete growth cycles. The growth of these parasites was monitored over 72 h with Giemsa-stained thin smear microscopy, counting at least 100 parasites per condition per timepoint, for 3 biological replicates. This experiment was repeated using NF54e parasites and Δ*Pf*MD3^1.1^ with and without the addition of 2 mM choline chloride to disentangle the contribution of sexual commitment to the multiplication rate of each strain. For this experiment, choline treated and untreated cultures at 0, 24, 48 and 72 h were fixed in 4% formaldehyde, 0.0075% glutaraldehyde in 1x phosphate buffered saline (PBS) at pH 7.4. Fixative was removed by washing samples 3x with 1x PBS prior to incubation with 1x SYBR Green I (Fisher Scientific, BMA50513) in PBS for 1 h shaking at 22 °C. Following staining, parasites were washed three times with PBS and parasitemia determined on an Attune Flow Cytometer, using SYBR Green I-stained uninfected erythrocytes as background control for gating. To calculate the multiplication rate, the parasitemia at 72 h was divided by the parasitemia measured at 24 h.

#### Sexual development phenotyping

To determine if there is a sexual growth phenotype that results from genetic disruption of *Pf*MD3, parasites from both clones (Δ*Pf*MD3^1.1^ and Δ*Pf*MD3^1.2^) and the NF54e parent strain were seeded as before but 48 h after seeding, gametocytogenesis was induced via nutrient starvation as described in ^7^ where, starting at a 1% parasitemia of majority ring stages in 4% hematocrit, 50% spent media was kept on the culture for 72 h. After the 72 h of nutrient starvation, the fresh RPMI media was supplemented with 20 U/ml heparin (Sigma) to prevent subsequent reinvasion of dividing asexual parasites^122^. For obtaining late-stage gametocytes, heparin treatment was continued for two full cycles (96 h) before culturing in normal supplemented RPMI media for the rest of the experiment. All cultures were maintained with daily medium changes and monitored with Giemsa-stained thin smear microscopy. This method of gametocyte induction and culturing was used for all experiments in this study except for studying gamete production or mosquito infection.

Growth phenotyping involved the calculation of % sexual conversion, percent gametocytemia and sex ratio. These characteristics were determined by Giemsa-stained microscopy at both 48 h (Stage II) and 144 h (Stage V) after the first addition of heparin to capture early and late-stage gametocyte development for three biological replicates and sex ratio of late stage gametocytes was determined by morphology using characteristics described in ^123^.

#### Mosquito infection for growth phenotyping

*Anopheles stephensi* infection with *P. falciparum* NF54e and the Δ*Pf*MD3 and Δ*Pf*MACFET mutants made in the NF54e parasite background was performed as previously described^124^. Asexual cultures were maintained *in vitro* in O+ erythrocytes at 4% hematocrit in RPMI 1640 (Corning) supplemented with 74 mM hypoxanthine (Sigma), 0.21% (wt/vol) sodium bicarbonate (Sigma), and 10% (vol/vol) heat-inactivated human serum (Interstate Blood Bank). Cultures were maintained at 37°C in a candle jar made from glass desiccators. Gametocyte cultures were initiated at 0.5% parasitemia and 4% hematocrit. Medium was changed daily for up to 15 to 18 days without the addition of fresh blood to promote gametocytogenesis. Cultures with final gametocytemias of 0.3% in 40% hematocrit containing fresh O+ human serum and O+ human erythrocytes were used for mosquito feeding. Similarly, for the genetic cross of Δ*Pf*MD3 and ΔMACFET, the final blood meal was composed of a 1:1 ratio of Δ*Pf*MD3 and ΔMACFET at 0.3% gametocytemia. *Anopheles stephensi* mosquitoes (3 to 7 days after emergence) were allowed to feed through a glass membrane feeder for up to 30 min on gametocyte cultures at 40% hematocrit containing fresh O+ human serum and O+ erythrocytes. Infected mosquitoes were maintained for up to 19 days at 25°C with 80% humidity and were provided with a 10% (wt/vol) sucrose solution.

#### Exflagellation and quantification of mosquito stage parasites for phenotyping

To assess the ability of each parasite line to exflagellate, mature gametocyte cultures at 0.5% gametocytemia (day 15 post-induction) were resuspended in 10% human serum and incubated at room temperature for 15 min as previously described^124^. The cellular suspension was placed on a hemocytometer and the number of exflagellating centers was counted using light microscopy at 40x magnification for three independent biological replicates. Samples were also collected for flow cytometric analysis at 0-, 10- and 15-min post-activation to measure the nuclear content over time. Parasites were fixed with 4% formaldehyde, 0.0075% glutaraldehyde in 1x PBS and stored at 4°C. For flow cytometry, fixative was removed by washing three times with PBS before incubation with 1x SYBR Green I (Fisher Scientific, BMA50513) in PBS for 1 h while shaking. Following staining, parasites were washed three times with PBS and nuclear content determined on an Attune Flow Cytometer, using SYBR Green I-stained uninfected erythrocytes as background control for gating. On day 12 post blood meal, mosquito midguts were dissected, and oocysts were stained with 0.05% mercurochrome in PBS. The midguts were then transferred to a 3-well slide containing 0.05% mercurochrome and oocysts were counted under 4x bright field microscopy. For salivary gland sporozoite counts, 15-25 salivary glands were dissected on day 14 or 15 post blood meal and pooled, homogenized, and sporozoites were counted using a hemocytometer. Two biological replicates were performed for the NF54e and Δ*Pf*MD3^1.1^ each and an additional replicate was performed on Δ*Pf*MD3^1.2^ alone and two biological replicates of Δ*Pf*MD3^1.1^ were crossed with ΔMACFET (a kind gift from Stefan Kappe and Sudhir Kumar, Seattle Children’s, Seattle, WA, USA). Graphs were created and statistical tests performed (either unpaired two-tailed t-tests or one-way ANOVA for Figure panel 2G) in Graphpad Prism v 10.1.0.

### Immunofluorescence assays and western blots

For immunofluorescence assays, parasites were fixed with 4% formaldehyde, 0.0075% glutaraldehyde in 1x PBS at pH 7.4 at 37°C for 45 mins followed by overnight incubation at 4°C. They were then washed three times in 1x PBS and permeabilized with 0.15% Triton X-100 for 10 mins at room temperature (RT). Potential left-over fixative was neutralized using 0.1 mg/mL NaBH_4_ for 4 mins at RT. Incubation with either rat anti-HA (Santa Cruz, sc-53516) or mouse anti-HA (Santa Cruz, sc-7392) and mouse anti-P230p^87^ (a kind gift from Dr. Kim Williamson, Uniformed Services University of the Health Sciences, Bethesda, MD, USA) or polyclonal rabbit anti-P47 (a kind gift from Dr. Alvaro Molina-Cruz, National Institutes of Health) primary antibodies (1:500, 1:1000 respectively) was carried out in 1% BSA overnight at 4°C. Subsequently, parasites were washed twice with PBS + 0.1% Tween 20, followed by two additional washes with PBS. Secondary antibodies conjugated to rat Alexa-488 (1:1000, Invitrogen, ab150157) or mouse Alexa-568 (1:500, Invitrogen, A10037) were prepared in 1% BSA and incubated overnight at 4°C. Parasites were washed twice with PBS + 0.1% Tween-20 and twice with PBS. The parasites were resuspended in SlowFade antifade (Invitrogen, S36973) containing DAPI according to manufacturer instructions and imaged on a Zeiss Axio Imager M2. Images were processed using Zen 2.6 pro with Fast Iterative deconvolution. For enumerating P230p and P47 positive and negative cells, slides were scanned with DIC to confirm presence of gametocytes and further verified to be DAPI positive prior to counting P230p or P47 positive and for the *Pf*MD3-3xHA line, HA-positive cells with at least 30 gametocytes counted per replicate with two biological replicates performed for each experiment. Results were graphed in Graphpad Prism v 10.1.0.

For western blotting, 0.05% saponin in PBS (v/v) was added to the parasite culture and incubated for 2 min to allow for lysis of RBC membranes. The resulting parasite pellet was washed 3 times with PBS and resuspended in lysis buffer (10 mM Potassium chloride (KCl), 20 mM HEPES (pH 7.0), 0.5% NP40, 1 mM dithiothreitol (DTT), 1x complete protease inhibitor cocktail (Sigma-Aldrich, 11836170001)) and incubated for 15 min at 4°C. The lysates were centrifuged at 2500 x *g* for 10 min at 4°C before the supernatant was collected. For cellular fractionation, the incubation was shortened to 10 min and two washes with lysis buffer preceded nuclear extraction by incubation in extraction buffer (800 mM Potassium chloride (KCl), 20 mM HEPES (pH 7.0), 1 mM DTT, 1x complete protease inhibitor cocktail) prior to SDS-PAGE and transfer to a nitrocellulose membrane in 1x Tris-glycine buffer with 10% methanol at 4°C. Membranes were blocked for at least 1 h in 5% milk and 1x TBS with 0.1% Tween 20 prior to incubation with primary antibody overnight at 4°C (anti-HA (Santa Cruz, sc-53516) 1:2500, anti-P230 (BEI Resources, MRA-878A) or anti-P47 1:1000, (anti-*Pf*aldolase (Abcam, ab207494)- 1:10000, anti-histone H3 (Abcam, ab18521) 1:10000) shown in Figure S4C. Secondary antibodies were added (anti-rat HRP (Abcam, ab7097) 1:2500, anti-mouse HRP (Invitrogen, A16017) 1:5000, anti-rabbit HRP (Abcam, ab6721) 1:10000) in 5% milk and TBS-0.1% Tween for 1 h at room temperature and developed using an ECL chemiluminescent detection kit (Thermofisher, 32209) prior to visualization on a Bio-Rad gel imager.

### Bulk RNA-sequencing for transcriptome analysis

Gametocyte cultures were prepared as described above. For the NF54e parent and Δ*Pf*MD3^1.1^ parasite lines, gametocytes were sampled on day 5 and 11 (n = 2) to capture early and late-stage gametocytes. Parasite samples were incubated with 0.05% saponin for 3 min and washed with PBS 3x before storage at -80°C until RNA preparation. RNA was prepared by phenol-chloroform extraction prior to using a Qiagen RNeasy kit (Qiagen, Germany, 74104) to purify the RNA as per manufacturer’s instructions. Integrity and quality of the RNA was confirmed by RNA Agilent TapeStation with RNA screen tape (5067-5576) prior to Illumina Truseq (MS-102-3001) RNA-sequencing library preparation.

RNA-sequencing results were analyzed using Galaxy v24.1.3 to perform pseudoalignment and relative quantifications (TPM) of reads using Kallisto v0.48^125^ against the PF3D7_genome version 59.0. Differentially expressed transcripts were identified using DEseq2 v 2.11.40.8^126^ on Galaxy with *p <* 0.05 and a Log_2_ Fold Change of Δ*Pf*MD3/NF54e in the 95^th^ percentile. Differentially expressed transcripts were clustered according to Euclidean distance in Rstudio v4.2.3 and visualized using the Pheatmap v1.0.12 package. Male and female transcripts were sourced from bulk RNA-seq data from separated male and female gametocytes in Lasonder *et al*.^9^ and single cell qPCR on male and female gametocytes from Walzer *et al.*^15^. For the bulk sequenced datasets, male and female differential transcripts were filtered for RPKM ≥20 in at least one sex, where the fold change between male and female transcripts was calculated and a cut-off of ± 2.0-fold change applied.

### Quantitative RT-PCR confirmation of transcript abundance in late-stage gametocytes

RNA samples used for transcriptomic analysis of NF54e parent and Δ*Pf*MD3^1.1^ parasite lines were treated with DNAse I (Sigma-Aldrich, AMPD1-1KT) for 30 minutes at 37°C. cDNA was synthesized from 500ng of total RNA using the RevertAid RT Reverse Transcription Kit (Thermo Scientific, K1691) according to manufacturer’s instructions. Gene expression was measured by reverse transcription followed by quantitative PCR (RT-qPCR) in biological duplicate, plated in technical duplicate, using Power SYBR™ Green PCR Master Mix (Applied Biosystems) on a CFX96 Touch Real-Time PCR Detection System (Bio-Rad). Gene primers used in the experiment can be found in Table S6. Primer sets used to profile P25 and P230p^127^, Pfs16 and 40S ribosomal protein S3^128^ using qPCR were previously published. Quantification of genes of interest was calculated by comparing each sample Cq value to a NF54e genomic DNA standard curve followed by normalization against sample 40S ribosomal protein S3 expression as adapted from ^129^ shown in Figure S4B.

### Single cell RNA-sequencing for transcriptome analysis

Δ*Pf*MD3^1.1^ parasites were maintained as described above. Gametocyte induction was initiated daily at the trophozoite stage (designated Day 0 for each culture) using 48 hours of conditioned medium stress^7^, generating temporally staggered cultures spanning Days 0–12 of development. Asexual stages (rings, trophozoites, and schizonts) and sexually committed parasites were represented by culture flasks corresponding to Days 0–2. From Day 2 onward, heparin (20 U/mL) was added to inhibit asexual replication and enrich Stage I–V gametocytes. Pooled asexual and sexual populations were developmentally offset by 24-hour intervals, enabling resolution of transcriptional transitions from commitment through early (Days 0–3), mid (Days 4–7), and late (Days 8–12) gametocyte stages.

Highly viable parasites were processed on the 10x Genomics Chromium platform for encapsulation into Gel bead-in-EMulsions (GEMs). Libraries were prepared using the Single Cell 3′ v3 kit post cDNA fragmentation and sequenced as technical duplicates on a NextSeq 2000 P2 platform to a depth of 30000 reads per cell^14^. Sequencing reads were mapped to the *P. falciparum* 3D7 genome (PlasmoDB v68.0) using Cell Ranger v9.0.1^130^.

Quality control and downstream analyses were performed in Seurat (v5.2.0)^131^. Cells with ≥1,000 UMIs and ≥300 detected genes were retained and predicted multiplets were removed using DoubletFinder (v2.0.4)^132^, yielding 7,026 high-confidence single cells spanning asexual and sexual developmental stages. Expression data were normalized, followed by dimensionality reduction and Louvain clustering to identify transcriptionally distinct cell populations, which were visualized using Uniform Manifold Approximation and Projection (UMAP)^133^.

To improve resolution of post-commitment sexual development and enable direct comparison with wild-type parasites, the *Pf*MD3-KO dataset was integrated with a published NF54 wild-type scRNA-seq dataset (Dogga *et al.*^14^) using SCTransform (v0.4.2)^134^ based integration with Harmony v1.2.3^135^ for batch correction. The integrated object was subsetted to retain sexual cells, including developing gametocytes, and sex-specific male and female clusters. Sub-clustering was performed to resolve finer developmental states, and clusters were annotated based on established stage- and sex-specific transcriptional markers^9,15^. Male and female signatures were further validated using module scoring in Seurat through AddModuleScore function with the top 100 sex-specific transcripts defined by Lasonder *et al.^9^*.

Developmental trajectories were inferred independently for male and female lineages using Slingshot (v2.14.0)^136^ to order cells along pseudotime. To identify genes with varying abundance dynamics influenced by loss of *Pf*MD3, generalized additive models were fitted along pseudotime using tradeSeq^137^ (genes ≥3 counts in ≥10 cells). Statistical significance (p < 0.001) was assessed using the FitGAM start, association, condition, and end tests to identify *Pf*MD3-KO effects on early gametocytogenesis, sex lineage progression, and terminal differentiation states. To identify biologically meaningful transcriptional changes, genes were required to exceed a Log₂ smoothed expression ≥ 0.5 in at least one condition (Δ*Pf*MD3 or NF54) within a lineage. Differential expression was evaluated using tradeSeq fitGAM models, integrating earlyDE, startVsEnd, diffEnd, and lineage-specific condition and association tests, with significance defined as padj < 0.001. Directionality was enforced using a minimum KO–WT difference of |Δlog₂| ≥ 0.5, while robustness of effects was further constrained by thresholds on mean Log_2_ Fold Change (≥ 1.0) and maximum absolute Log_2_ Fold Change (≥ 2.0).

### Proteomics of NF54e and *Pf*MD3 KO parasites

*P. falciparum* NF54e and Δ*Pf*MD3^1.1^ gametocytes were cultured as above until day 5 (Stage II-III) and day 11 (Stage V), lysed with saponin in PBS (0.05% v/v) and washed three times with PBS before being snap frozen and stored at -80°C until preparation for mass spectrometry. Samples were collected and analyzed in biological replicate (n = 2).

#### Proteomics sample preparation

Unless otherwise noted, all solid reagents were from Sigma-Aldrich; solvents and acids were Optima LC-MS grade from Fisher Scientific. Water was LC-MS grade from Honeywell Burdick & Jackson (LC365). To each frozen cell pellet was added 2x lysis buffer (10% sodium dodecyl sulfate (SDS) in 100 mM ammonium bicarbonate (NH_4_HCO_3_)). Samples were incubated 5 min at 95°C in a thermomixer at 1000 RPM, then centrifuged 1 min at 20,000 x *g* to settle the insoluble material. Water was added to adjust the buffer concentration to 1x, and the pellet was resuspended by pipetting. The sample was incubated for 5 more minutes at 95°C and 1000 RPM, then centrifuged twice for 1 min at 20,000 x *g*, rotating the tube 180° in between to settle the insoluble material. Protein concentration was measured by BCA assay. For each sample, an aliquot containing 20 µg total protein was added to a 2.0 mL microcentrifuge tube (Eppendorf Protein LoBind, 0030122283) and adjusted to a final volume of 60 µL with 1x lysis buffer.

To each sample was added 2 µL of 150 mM tris(2-carboxyethyl) phosphine (TCEP; Pierce Bond Breaker 77720) in 70 mM NH_4_HCO_3_. After incubating 15 min at 55°C, the samples were cooled to room temperature and 2 µL of 0.6 M iodoacetamide in 50 mM NH_4_HCO_3_ was added. The samples were incubated 20 min in darkness at room temperature in a thermomixer at 1000 RPM, then an additional 32 µL of 50 mM NH_4_HCO_3_ was added (final volume 96 µL). Samples were digested using SP3 protein aggregate capture^138^. Two different types of magnetic carboxylate beads (Cytiva, 24152105050250 and 65152105050250) were mixed 1:1, washed 3 times with 50 mM NH_4_HCO_3_, and resuspended at 50 mg/mL in 50 mM NH_4_HCO_3_ before adding 4 µL of beads (200 µg) to each sample. The samples were briefly vortexed to suspend the beads, and after adding 100 µL of acetonitrile (ACN), were incubated 20 min at room temperature on a thermomixer at 700 RPM. The supernatant was removed, and the beads were washed three times with 180 µL of 80% ACN. To each sample 1.0 µg of trypsin (platinum mass spectrometry grade, Promega, VA9000) was added in 40 µL of 100 mM NH_4_HCO_3_ and samples were incubated overnight at 37°C in a thermomixer at 700 RPM. After digestion, 40 µL of 1% TFA was added, and the samples were briefly vortexed to mix. The supernatants were recovered, transferred to clean 1.5 mL Protein LoBind tubes, pelleted 1 min at 20,000 x *g* to settle any residual beads, and transferred to autosampler tubes.

#### Liquid Chromatography (LC) with Data Independent Acquisition-Mass Spectrometry (DIA-MS)

LC was performed with an EASY-nLC 1000 (Thermo Fisher Scientific) using a vented trap set-up. The trap column was a PepMap 100 C18 (Thermo Fisher Scientific, 164946) with 75 µm i.d. and a 2 cm bed of 3µm 100 Å C18. The analytical column was an EASY-Spray column (ThermoFisher Scientific, ES904) with 75 µm i.d. and a 15 cm bed of 2µm 100 Å C18 operated at 35°C. The LC mobile phases (Honeywell Burdick & Jackson, LC452) consisted of buffer A (0.1% v/v formic acid in water) and buffer B (0.1% v/v formic acid in ACN). The separation gradient, operated at 300 nL/min, was 4% B to 28% B over 85 min, 28% B to 30% B over 5 min, 30% B to 80% B over 5 min, and 10 min at 80% B. Prior to each run, the trap was pre-conditioned with 12 µL buffer A at 800 bar, the column was preconditioned with 6 µL buffer A at 800 bar, and the sample was loaded onto the trap with buffer A. Data-independent acquisition mass spectrometry (DIA-MS) was performed with a Thermo Fisher Scientific Orbitrap Eclipse. The samples were analyzed with the following acquisition settings: MS1 scan from 395-1005 m/z at 30,000 resolution, default AGC settings (target of 4×10^5^ ions, max ion injection time set to “Auto”); MS2 at 15,000 resolution with an AGC target of 1000% (5×10^5^ ions) and a maximum injection time of 22 ms, HCD fragmentation at 33% normalized collision energy, 8 *m*/*z* isolation window, and a loop count of 75 with an inclusion list of 151 overlapping DIA windows with optimized *m*/*z* values generated using the tool in EncyclopeDIA version 1.12.31. In order to produce a spectral library, the eight samples were combined into single pooled sample and analyzed using gas-phase fractionation^139^, a method wherein the same sample was injected six times with each method narrowed to a 100 *m/z*-wide band of masses. The DIA-MS methods were the same as above except for the following: MS1 scan range was narrowed to 100 *m/z*; MS2 maximum injection time of 54 ms, 4 *m/z* isolation window, and a loop count of 25 with an inclusion list of 51 overlapping DIA windows.

#### DIA data analysis

Raw mass spectrometry data were converted to mzML using msConvert version 3.0.19106 (Proteowizard^140^). Peaks were centroided and the staggered DIA windows were demultiplexed^141^. DIA data were analyzed with EncyclopeDIA version 1.12.31^141^ using Open JDK 13 on a Slurm 19.05.5 cluster running under Ubuntu 20.04. EncyclopeDIA analyses were executed in the command line using default parameters except that Percolator version 3-01 was specified and the Percolator training set size was increased to 1M. A spectral library was generated *in silico* using Prosit^142^ as described previously^141^. Briefly: a protein FASTA database was assembled from the reference *P. falciparum* 3D7^143^ database (www.PlasmoDB.org^99^ version 59); a human database (Swiss-Prot reviewed reference proteome obtained from UniProt.org^144^ the common Repository of Adventitious Proteins (www.thegpm.org/cRAP); and the selection marker BSD. The sequences were digested *in silico* using the tool in EncyclopeDIA^145^ to obtain a list of all theoretical peptides with *m/z* between 396.4 and 1004.7, two tryptic termini, up to one missed cleavage, and charge state 2 or 3. This list was uploaded to Prosit (www.proteomicsdb.org/prosit) and theoretical spectra were generated for the peptides using the 2020 HCD intensity prediction model and the 2019 iRT retention time prediction model, assuming an NCE of 33 and default charge state of 3. The pooled library sample (comprising six gas-phase fractions) were searched against this *in silico* library. Using tools in EncyclopeDIA, the results of these searches were combined and exported as a chromatogram library comprising the best peptide spectra and retention times empirically derived from pooled sample under the same LC-MS conditions used to collect the individual sample data. The DIA-MS data from the samples was searched against this empirical chromatogram library. Protein quantification, including match-between-runs, was performed using default settings. Protein peak areas for any protein in each sample were replaced with zero if no peptides matching that protein were detected in that sample. Raw protein peak areas were quantile normalized for detected *P. falciparum* protein areas in each sample. Results were visualized in a volcano plot using ggplot2 v3.5.1 and Pheatmap v1.0.12 in Rstudio v4.2.3 with lines indicating differentially abundant proteins with Log_2_FC ≥2 and *p-*value <.05. In cases where proteins were not detected in a specific sample, the minimum value detected in that sample was substituted for the calculation of Log_2_ fold changes and *P*-value was set to 1 to indicate that the error in the observation could not be determined.

### RNA Bind-N-Seq and RNA electrophoretic mobility shift (EMSA) *in vitro* binding assays

For *in vitro* RNA-binding assays, the full length, short (aa 140-214), medium (aa 1-214) and long (aa 140-485) versions of *Pf*MD3 were cloned downstream of a GST-tag in the pGEX6P1 (Cytiva, 28-9546-48) vector for recombinant protein expression in bacteria, *E. coli* BL21(DE3)pLysS (Promega, L1195). The full-length protein was not detectably expressed under any tested conditions, and the long and short constructs were used for subsequent experiments. For protein expression, bacteria were grown from a starter culture in 100 or 500 mL LB broth with 100 µg/mL ampicillin and 35 µg/mL chloramphenicol at 37°C until OD_600_ of approximately 0.6 before induction with 0.2 mM IPTG. Following induction, the temperature was reduced to 25°C and 0.1 mM zinc chloride was added to bacterial culture and protein was expressed for 2 h before freezing the bacterial pellet at -80°C until lysis and protein purification. Prior to purification, bacteria were lysed in BPER reagent (Thermo Fisher Scientific, 89821) at room temperature for 10 min. Purification of the GST-tagged proteins was performed using a GST Fusion Protein purification kit (Genscript, L00207) per the manufacturer’s instruction with the exception that the pH of the gravity flow buffer was decreased to 6.8 to prevent chelation of the zinc from the CCCH-type zinc finger in the *Pf*MD3 protein^146^ before elution at pH 8.0 and 10 U of DNAse (Biobasic, BS88253) and 10 U RNAse A (Biobasic, RB0473) were added to bacterial lysate.

<u>R</u>NA <u>B</u>ind-<u>N</u>-<u>S</u>eq (RBNS) was carried out as described in^93^, using 50, 100, 250 and 500 nM concentrations of the short *Pf*MD3 construct and a 40-mer random RNA library transcribed *in vitro*. The same concentrations of proteins were used to test *Pf*MD3 in the presence of 5mM EDTA. The GST-*Pf*MD3-RNA complex was pulled down using glutathione high-capacity magnetic agarose beads (Sigma-Aldrich, G0924). Following RBNS, the *Pf*MD3 bound RNA was purified on Zymo RNA Clean and Concentrate columns (R1016) and eluted into nuclease-free water prior to reverse transcription and sequencing library preparation via PCR with primers and indexes for Illumina Truseq RNA libraries. The input library control was reverse transcribed and amplified alongside the immunoprecipitated samples. RBNS libraries were pooled and single-end sequencing performed with a Miseq V3 at 10 pM concentrations. Data from RBNS were analyzed in Linux using scripts from the C. Burge lab RBNS pipeline (https://github.com/cburgelab/RBNS_pipeline) and the most enriched motifs relative to the sequenced input library reported.

For RNA <u>E</u>lectrophoretic <u>M</u>obility <u>S</u>hift <u>A</u>ssays (EMSAs) the short *Pf*MD3 protein purified as described above was incubated with a 5’ biotinylated probe with sequence 5’-GUAGG<u>GAAGAA</u>G<u>GAA</u>AGG-3’ corresponding to the region under the CLIP-seq peak observed in the *p230* gene body or a scrambled version (https://www.bioinformatics.org/sms2/shuffle_dna.html) of this same sequence 5’-GGGA<u>GAA</u>AGAUGAGGAGA-3’. An additional probe was included as a negative control that matched the *p230* sequence but had the GAA regions mutated to CUU residues 5’-GUAGG<u>CUUCUU</u>G<u>CUU</u>AGG-3’. Competitive binding was measured in each reaction mixture by the addition of 10, 100 or 1000-fold molar excess of unlabeled RNA probe to the included biotinylated probe. RNA EMSAs were performed as per manufacturer’s instructions for the Thermo Scientific LightShift Chemiluminescent RNA EMSA Kit (20158) using 5 nM of the short *Pf*MD3 containing the CCCH-type zinc finger domain (aa 140-214). Gel images captured using a BioRad gel imager with Auto-optimal detection of chemiluminescence set for 3x3 captures.

### Immunoprecipitation-Mass Spectrometry (IP-MS) and data analysis

*Pf*MD3 3xHA tagged parasites were grown in biological replicate (n = 3) to early-stage gametocytes (day 5 post-induction) and samples collected by centrifugation. Pelleted parasite cultures were resuspended in regular RPMI prior to adding 0.05% saponin in PBS (v/v) and incubating for 2 min at 4°C to allow for lysis of RBC membranes. The lysed parasite pellet was washed 3x in PBS and passed through a 22 g needle before being resuspended in lysis buffer (100 mM Potassium chloride (KCl), 5 mM magnesium chloride (MgCl_2_), 10 mM HEPES (pH 7.0), 1 mM EDTA, 0.5% NP40, 1 mM DTT, 1x complete protease inhibitor cocktail), passed through a 22g needle again and incubated at 4°C for 15 min with rotation to release parasite proteins. The lysed parasites were centrifuged at 2500 x *g* for 10 min at 4°C before the supernatant was added to 1/4^th^ volume of dilution buffer (40% glycerol, 20 mM HEPES (pH 7.0), 1 mM EDTA, 1 mM DTT, 1x complete protease inhibitor cocktail) and 10 µg α-HA antibody (Roche, 3F10) or 20 µg IgG antibody (Millipore Sigma, PP68) coupled to M270 epoxy Dynabeads (Thermo Fisher Scientific, 14301) per the manufacturer’s instructions. Proteins were Immuno<u>p</u>recipitated (IP) by rotation for 2 h at 4°C before 3x washes with dilution buffer without glycerol, transfer to a tube and 3x washes in PBS with a second tube transfer. The PBS was removed from the beads as completely as possible before storage at -80°C until the samples could be processed with DIA-MS.

#### DIA-MS sample processing and data analysis

For each sample, 0.4 µg of trypsin in 20 µl 50 mM NH_4_HCO_3_ was added to washed and ready to digest beads and incubated at 37°C overnight. The supernatant was separated from the magnetic beads and pH was adjusted to 3 with 10% formic acid. The sample was desalted with stage tip cleanup before LC-MS. Samples were analyzed by LC-MS using Nano LC-MS (Dionex Ultimate 3000 RLSCnano System, Thermofisher) interfaced with Eclipse (Thermofisher).

Samples were loaded on to a fused silica trap column Acclaim PepMap 100, 75 µm x 2 cm (ThermoFisher). After washing for 5 min at 5 µl/min with 0.1% TFA, the trap column was brought in-line with an analytical column (Nanoease MZ peptide BEH C18, 130A, 1.7 µm, 75 µm x 250 mm, Waters) for LC-MS/MS. Peptides were fractionated at 300 nL/min using a segmented linear gradient 4-15% B for 30 min (where A: 0.2% formic acid, and B: 0.16% formic acid, 80% ACN), 15-25% B for 40 min, 25-50% B for 44 min, and 50-90% B for 11 min. Solution B then returns to 4% for 5 minutes for the next run. A DIA (Data independent acquisition) workflow was used to analyze the eluted peptides. MS scan range were set to 400-1200, resolution 12,000 with AGC set at 3E6 and ion time set as auto. 8 m/z windows were set to sequentially isolate (AGC 4E5 and ion time set at auto) and fragment the ions in C-trap with relative collision energy of 30. The MS were recorded with Resolution of 30,000. Raw data were analyzed with predicted library from “P.falciparum3D7.fasta” database for library-free search using DIA-NN 1.8.1^147^ with recommended settings. The results were filtered for both PEP (an estimate of the posterior error probability for the precursor identification, based on scoring with neural networks) filtered <0.01 and PG.Q (Protein Group Q Value) filter <0.01. Protein group MaxLFQ value were used for group comparisons. Enriched proteins in the IP were defined as those >2x enriched in the IP of *Pf*MD3-3xHA compared to IgG in all three biological replicates. Results were visualized using Microsoft Excel and, in cases where proteins were not detected in a specific sample, the minimum value detected in that sample was substituted to the fold change calculation of the IP with HA compared to the IgG negative control. GO-term enrichments were performed and visualized using ShinyGO 0.80 (https://bioinformatics.sdstate.edu/go) and filtered for FDR <0.05, showing the top 10 pathways with a minimum number of two genes per pathway and removing redundant pathways from the results.

### Crosslinked Immunoprecipitation RNA-sequencing (CLIP-seq) and data analysis

*Pf*MD3 3xHA tagged parasite cultures were induced to gametocytogenesis as described above. On day 5 or 11 post-induction, the heparin containing media was removed by centrifugation and replaced with normal RPMI media before adding 0.05% saponin in PBS (v/v) and incubating for 2 min to allow for lysis of RBC membranes. The resulting parasite pellet was washed three times with PBS before being passed through a 22g needle into a sterile petri dish. The parasites were then crosslinked under UV light at 254 nm to a total energy of 400 mJ. The parasites were centrifuged and resuspended in lysis buffer (100 mM Potassium chloride (KCl), 5 mM magnesium chloride (MgCl_2_), 10 mM HEPES (pH 7.0), 0.5 % NP40, 1 mM DTT, 20 U/mL Superasin RNAse inhibitor (AM2696), 1x complete protease inhibitor cocktail. The parasites were again passed through a 22g needle before incubating for 15 min on ice. The lysed parasites were centrifuged at 2500 x *g* for 10 min at 4°C before the supernatant was added to 10 µg α-HA antibody (Roche 3F10) or 20 µg IgG antibody (Millipore, PP68) coupled to M270 epoxy Dynabeads (Thermo Fisher Scientific,14301). Before adding the antibodies, 1% of the lysate was removed and kept as an input control. The samples were rotated at 20 rpm at 4°C for 2 h before collecting the antibody-coupled beads on a magnetic stand and performing washes three times with wash buffer (50 mM Tris–HCl (pH 7.4), 150 mM NaCl, 1 mM MgCl_2_, 0.05 % NP40). Next, wash buffer supplemented with 30 µg proteinase K was added to each sample and incubated at 55°C for 20 min. The RNA was released from the RNP complexes by the addition of 4x volumes Trizol reagent (Thermo Fisher Scientific, 15596018). RNA was extracted and purified as for the RNA-seq before Illumina Truseq total RNA-sequencing library preparation (MS-102-3001).

Analysis of the sequenced reads obtained from CLIP-seq was conducted on using CLIP-explorer Galaxy Project (https://clipseq.usegalaxy.eu/)^148^. Following assessment of the quality of the reads using FastQC v 0.74, adapter sequences were trimmed using Trimmomatic v 0.39^149^ and reads mapped to the *P. falciparum* 3D7 genome (Pf3D7 v59, obtained from www.PlasmoDB.org^99^) using BWA MEM2 v2.2.1^118^. Reads were filtered using SAMtools to remove duplicates. Peaks were called on the IP and IgG samples relative to the input control using PureCLIP^150^ for each replicate. Following peak calling, peaks in the IP were retained if they obtained at least 2-fold enrichment over IgG samples and were present in at least two replicates of the IP and read coverage over peak regions was >10 reads. Additional peaks were called using Piranha^151^ where bigwig files were generated by subtracting the input control from the HA-IP and IgG-control samples using bamcompare. These bigwig files were then used as input for Piranha, where the IgG control was specified as the covariate, reads were binned into 100 bp regions, and peaks were called with a *P-*value of <0.01 for the IP relative to the IgG control. Peaks called in Piranha were retained if they were called in at least two biological replicates and at least 10 reads were mapped across the region in the IP sample. Peaks were visualized in IGV v2.13.0^152^ and figures were created in Graphpad Prism v10.1.0. Transcript FASTA sequences from RNA targets were searched for motif enrichment using STREME (https://meme-suite.org/meme/doc/streme.html^92^) using the *P. falciparum* 3D7 transcriptome (Pf3D7 v59, obtained from www.PlasmoDB.org^99^) as background.

### Assessment of translation efficiency with *in vitro* translation assay

Translation-competent lysate was generated from stage II gametocytes from NF54e or Δ*Pf*MD3 parasites as described previously ^153^. These parasite lysates were incubated with 1x reaction mix and 2.5 µl accessory proteins from a 1-Step human coupled *in vitro* translation (IVT) kit (Thermo Scientific, 88881) for 2 h at 37°C. As an additional condition, Δ*Pf*MD3 lysate was incubated in a IVT reaction as described but also including 2 µg of the long version of recombinant *Pf*MD3-GST. Before incubation, a baseline measurement of protein concentration within each reaction was carried out using a BCA protein assay (Fisher Scientific, PI23227) with measurements compared against a bovine serum albumin (BSA) standard curve. Following incubation, the endpoint protein concentrations were determined with the BCA assay.

### Confirmation of protein-protein interactions from IP-MS with protein pull-down

*Pf*MD3 3xHA tagged parasite lysate was prepared using the same method described for IP-MS for two biological replicates. Two protein interactors were selected for validation, MPP10 (PF3D7_1027100) and AMP-deaminase (PF3D7_1329400). For each gene, the full coding sequence was codon-optimized for *E. coli* and cloned into pET28a expression vectors and the full construct purchased from Twist Biosciences. The plasmids were transformed into Rosetta pLysSDE3 competent cells (EMD Millipore, cat. no. 71401) and inoculated from a starter culture into 200 mL LB broth with 50 µg/mL kanamycin and 35 µg/mL chloramphenicol at 37°C until OD_600_ of approximately 0.6 before induction with 0.2 mM IPTG. Protein expression proceeded at 30°C for 5 h prior to harvesting. Prior to purification, bacteria were lysed in BPER reagent (Thermo Fisher Scientific, cat. no. 89821) at room temperature for 10 min. Soluble proteins were collected by centrifugation at 12000 x *g* for 10 min at 4°C. The supernatant was combined with 2x volumes of 1x PBS with 10 mM imidazole, 1x complete protease inhibitor cocktail, 1 U/ml of DNAse and 1 U/ml RNAse A. The His-tagged proteins were purified using Pierce HisPur Affinity Purification Kits (Thermo Scientific, 88224) according to the manufacturer’s instructions and the proteins dialyzed using Amicon Centrifugal filters (Thomas Scientific, 1211M37). For the protein pulldown, *Pf*MD3 3xHA lysate from two biological replicates was combined with 20 µg of each His-tagged recombinant protein and mixed by gentle rotation for 30 min at 4°C. Next, 20 µl of magnetic Dynabeads for His-tag Isolation and pulldown (Fisher Scientific, 10103D) was added to each reaction, including a negative control where no recombinant protein was added. Following IP, beads were washed as described for IP-MS and beads were boiled at 95°C for 10 min in 4x LDS sample buffer (VWR, 76286-030) with 2% β-mercaptoethanol. Protein pull-down was confirmed via western blot with conditions as described above and probed for IP with 1:10 000 6xHis-HRP antibody (Abcam, ab237339).

### Determination of *Pf*MD3-HA association with translational machinery through sucrose gradient profiling

*Pf*MD3 3xHA tagged parasite lysate from stage II gametocytes was prepared for sucrose concentration as described previously^154,155^. The parasite lysate was clarified by centrifugation at 20 000 *x g* for 10 min at 4°C and loaded on a 35% (w/v) sucrose cushion in polysome buffer (400 mM potassium acetate, 25 mM potassium HEPES pH 7.2, 15 mM magnesium acetate, 200 μM cycloheximide (Sigma Aldrich, 239763), 1 mM dithiothreitol (DTT), and 1x cOmplete Protease Inhibitor Cocktail without EDTA, SuperAsin 1 µl/ml). The ribosomes were concentrated through the sucrose cushion by ultracentrifugation at 150 000 *x g* at 4°C for 2 h in a Beckman Coulter Optima centrifuge with a T100.3 rotor. Next, the ribosome pellets were resuspended in polysome buffer prior to layering onto a 2 mL continuous linear 15% to 60% (w/v) sucrose gradient and centrifuged at 200,000 *x g* at 4°C for 3 h in a TLS-55 swinging bucket rotor. Fractions of 100 µl were collected and absorbance at 254 nm of each fraction determined via UV spectrophotometer. Proteins were precipitated using 10% TCA (v/v) overnight at 4°C and centrifuged at 10,000 *x g* at 4°C for 20 min. Precipitated proteins were washed twice with ice cold acetone and air dried. Fractions were then boiled in 4x LDS loading dye with 2% β-mercaptoethanol prior to western blot analysis.

