## supplementary figures for "The CCCH-type zinc-finger *Pf*MD3 promotes translation for malaria parasite transmission"

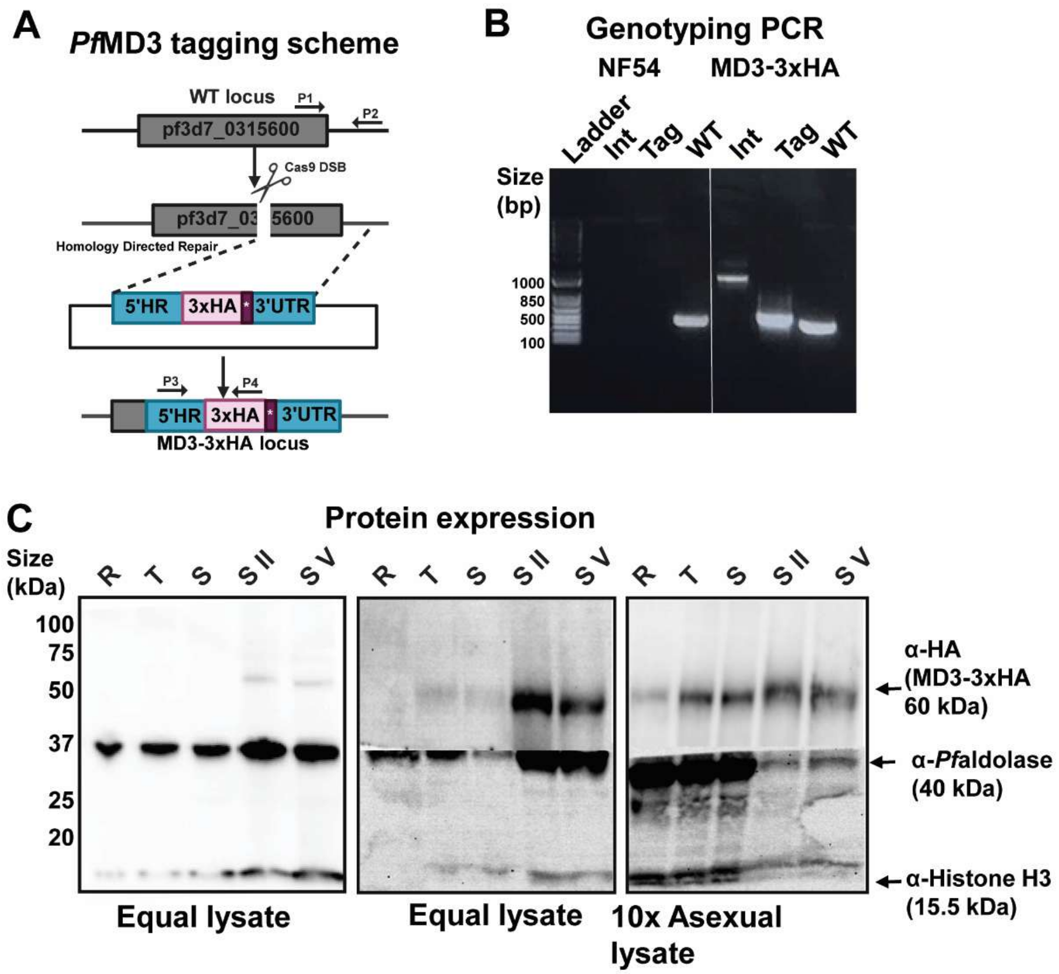


**Figure S1. Verification and expression of *Pf*MD3-3xHA, related to Figure 1.** (**A**) A CRISPR-Cas9 based approach[115] was used to produce a clonal *Pf*MD3-3xHA tagged line in NF54e parasites (MRA-1000A). (**B**) Primers P1+P2 (Table S6) were used as a positive control for the *Pf*MD3 locus (WT, 485 bp) while plasmid uptake was verified with P3+P4 (Tag, 612 bp) and integration of the tag at the 3' end of *Pf*MD3 verified with P1+P4 (**I**nt, 1095 bp). (**C**) Protein expression of *Pf*MD3 was verified by Western blot showing the uncropped membranes quantified in Figure 1D here, with signal intensity compared to the *Pf*aldolase control quantified by densitometry. Abbreviations: R = ring-stage, T = Trophozoite, S = Schizont, S **II** = gametocyte stage **II**, S V = gametocyte stage V, C = cytoplasmic extract and N = nuclear extract. Different lengths of exposure were applied to membranes for visualizing expression in asexual stages and gametocytes in order to obtain a clear image.

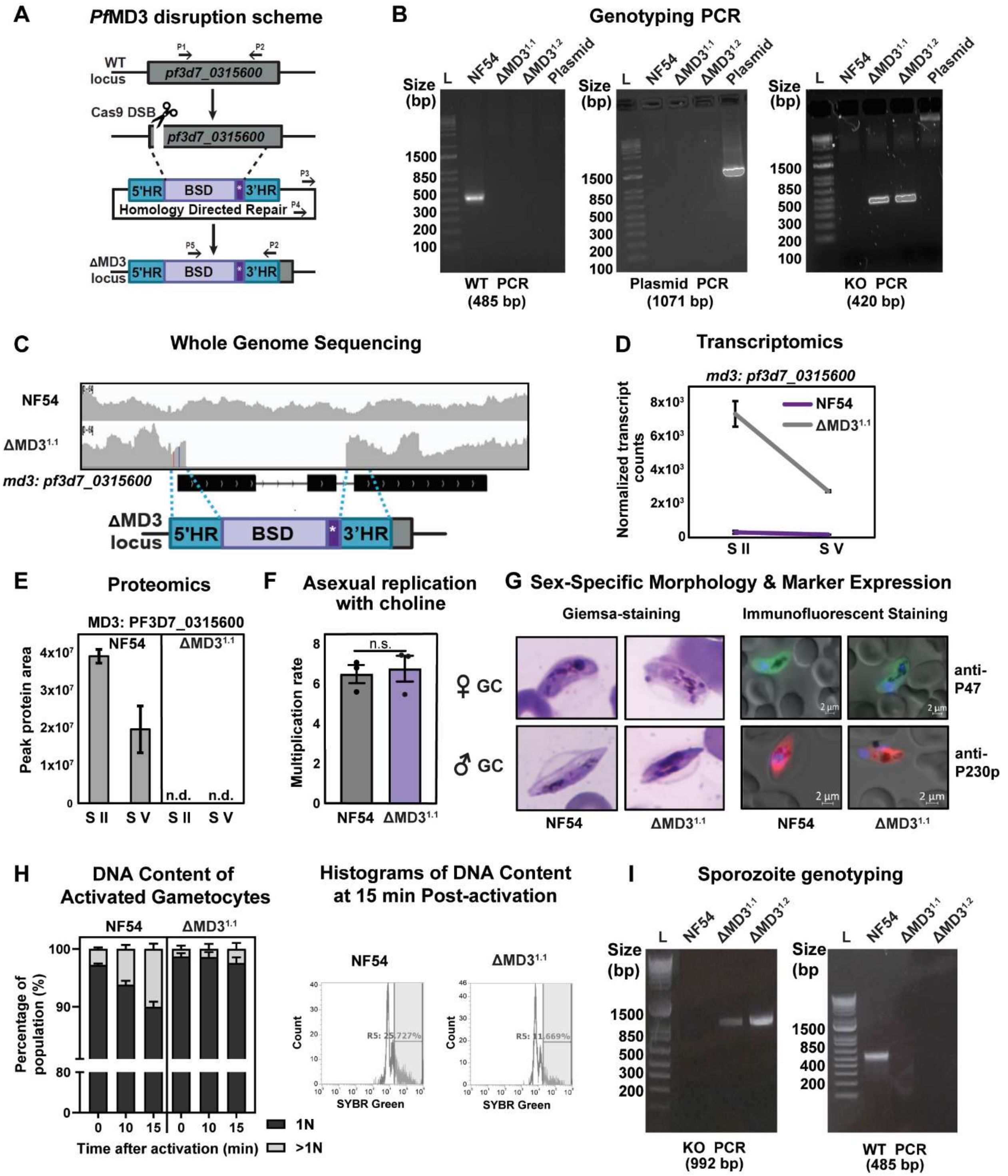


**Figure S2. Verification of *Pf*MD3 genetic knockout, related to Figure 2.** (**A**) Knockout strategy and (**B**) genotyping PCR verification of the NF54e parent (WT PCR, P1+P2, *Pf*MD3$^{1.1,1.2}$ (KO PCR, P5+P2), and the CRISPR Cas9 plasmid (P3+P4) by replacement of the 1$^{st}$ two exons of *Pf*MD3 with a blasticidin (BSD) resistance cassette following a Cas9 mediated double stranded break (DSB). (**C**) IGV screenshot of the MD3 locus from whole genome sequencing of the NF54e parental line and the Δ*Pf*MD3$^{1.1}$. Verification of the expression of the *Pfmd3* transcript in both the Δ*Pf*MD3$^{1.1}$ and NF54e parent strain from (**D**) next-generation RNA-sequencing and (**E**) DIA-MS throughout gametocyte maturation (n = 2). (**F**) Relative multiplication rate of Δ*Pf*MD3[1.1] and NF54e parent strain was determined in the presence of 2 mM choline chloride to prevent conversion to gametocytes in the culture (n = 3). (**G**) Representative morphology of mature stage IV-V gametocytes was captured for the WT NF54e parent compared to Δ*Pf*MD3[1.1] using light microscopy imaging of Giemsa-stained thin blood smears and immunofluorescent microscopy using either P230p antibody with mouse Alexa-568 secondary antibody to visualize male gametocytes or P47 antibody with rabbit Alexa-488 secondary antibody to visualize female gametocytes. (**H**) The ability of ΔMD3 to produce male gametes was investigated by inducing exflagellation of gametocytes (day 15) and measuring nuclear content (with example gating strategy next to graph) via flow cytometry following staining with SYBR Green I in technical triplicate. (**I**) PCR confirmation that sporozoites generated from the NF54 parent and Δ*Pf*MD3[1.1] parasites to the mosquito, retained either the wild-type or disrupted MD3 locus (Primers in Table S6).

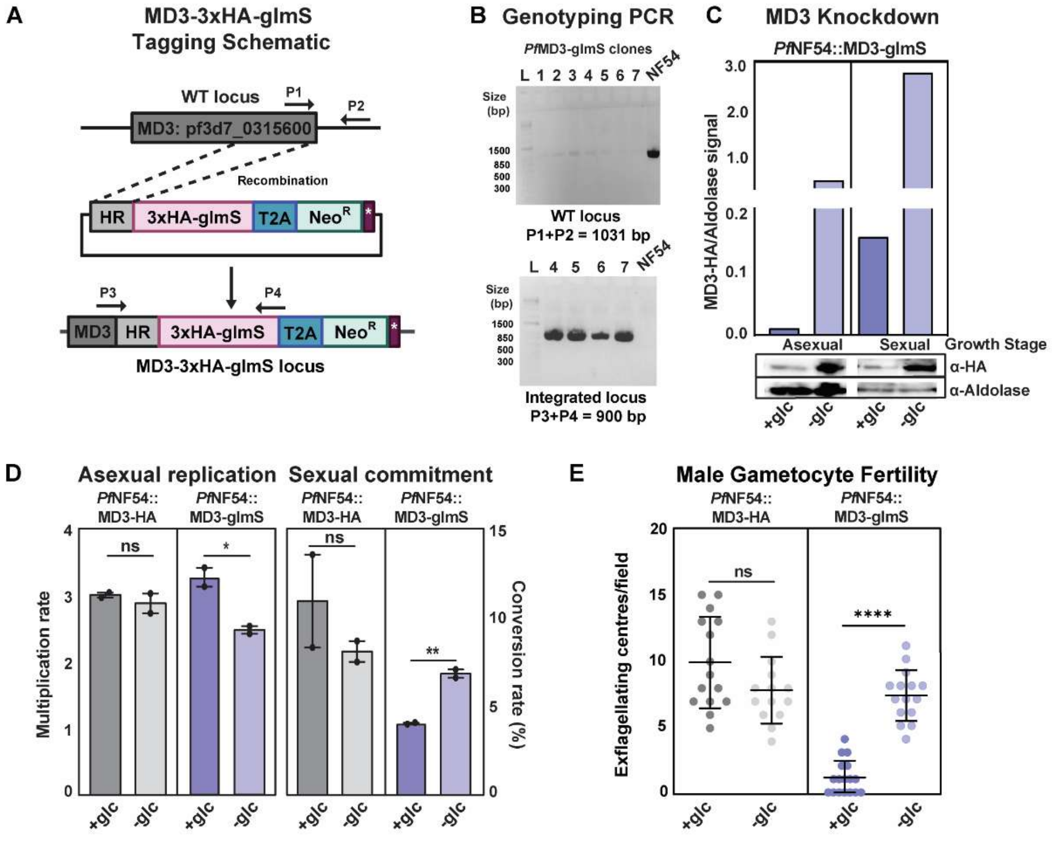


**Figure S3. Confirmation of the role of *Pf*MD3 in maintaining *P. falciparum* male fertility.** (**A**) The pSLI-glmS plasmid was used to produce a clonal line of *Pf*MD3-3xHA-glmS that would allow for glmS-ribozyme mediated knockdown of *Pf*MD3 mRNA upon addition of glucosamine. (**B**) PCR verification of PfMD3-3xHA-glmS clones (1-7) with NF54e parent strain control, clone 7 was used for follow up experiments. (**C**) Bargraph shows the degree of protein knockdown measured by western blot (HA signal compared to *Pf*aldolase) obtained for *Pf*MD3-3xHA-glmS following treatment with 1.25 mM glucosamine starting at either 8-12 hpi in asexual development or starting at Stage I of gametocytogenesis sexual parasites for 72 h. Following treatment with glucosamine, *Pf*MD3-3xHA-glmS or *Pf*MD3-3xHA parasites were assessed for (**D**) asexual blood stage replication rate, percent gametocyte conversion rate, or (**E**) The ability of to produce male gametes. Each experiment was conducted with n = 2 biological replicates and for panels B and C parasitemia was counted using flow cytometry with at least 10, 000 cells counted per sample while for D, exflagellation of gametocytes was induced on day 15 the number of exflagellating centers per field were counted on a hemocytometer.

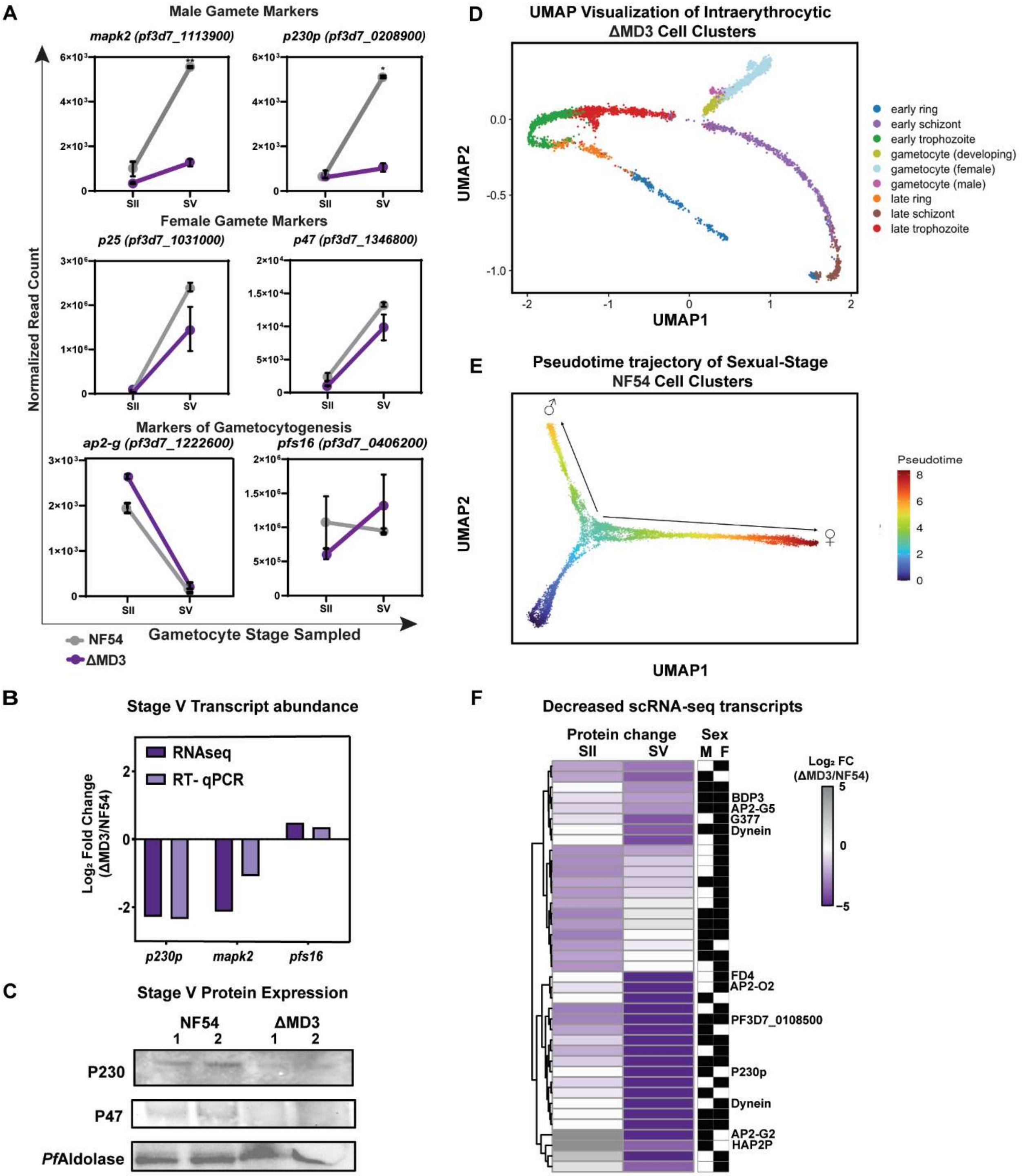


**Figure S4. Molecular profile of Δ*Pf*MD3 gametocytes, related to Figure 3.** (**A**) Known markers of gametocytogenesis and male or female development are plotted as line graphs with S.E.M. between the biological replicates of the WT and KO lines respectively (n = 2) and significant differences from WT are indicated (Two-tailed t-test, * = $p < .05$, ** = $p < .01$). (Associated data in Table S2). (**B**) Comparison of transcript abundance of male markers P230p, MAPK2 and Pfs16 and P25 gametocyte markers in Δ*Pf*MD3 compared for NF54 as measured by RT-qPCR and RNA-seq respectively. (**C**) Validation of differential expression of P230 and P47 expression from proteomics data with western blot for n = 2 biological replicates (1,2) as compared to *Pf*aldolase loading control. (**D**) UMAP trajectories of all *Pf*MD3$^{1.1}$ cells integrated into the Dogga et al.[14] UMAP trajectories for NF54. (**E**) Slingshot-based lineage and pseudotime inference overlaid on the UMAP embedding of 3,106 subsetted ΔPfMD3$^{1.1}$ cells integrated with 7,988 subsetted NF54 gametocyte cells from the Malaria Cell Atlas. (**F**) Heatmap representation of differentially abundant scRNA-seq transcripts from either the male or female lineage (M,F) against stage II and stage V proteomics.

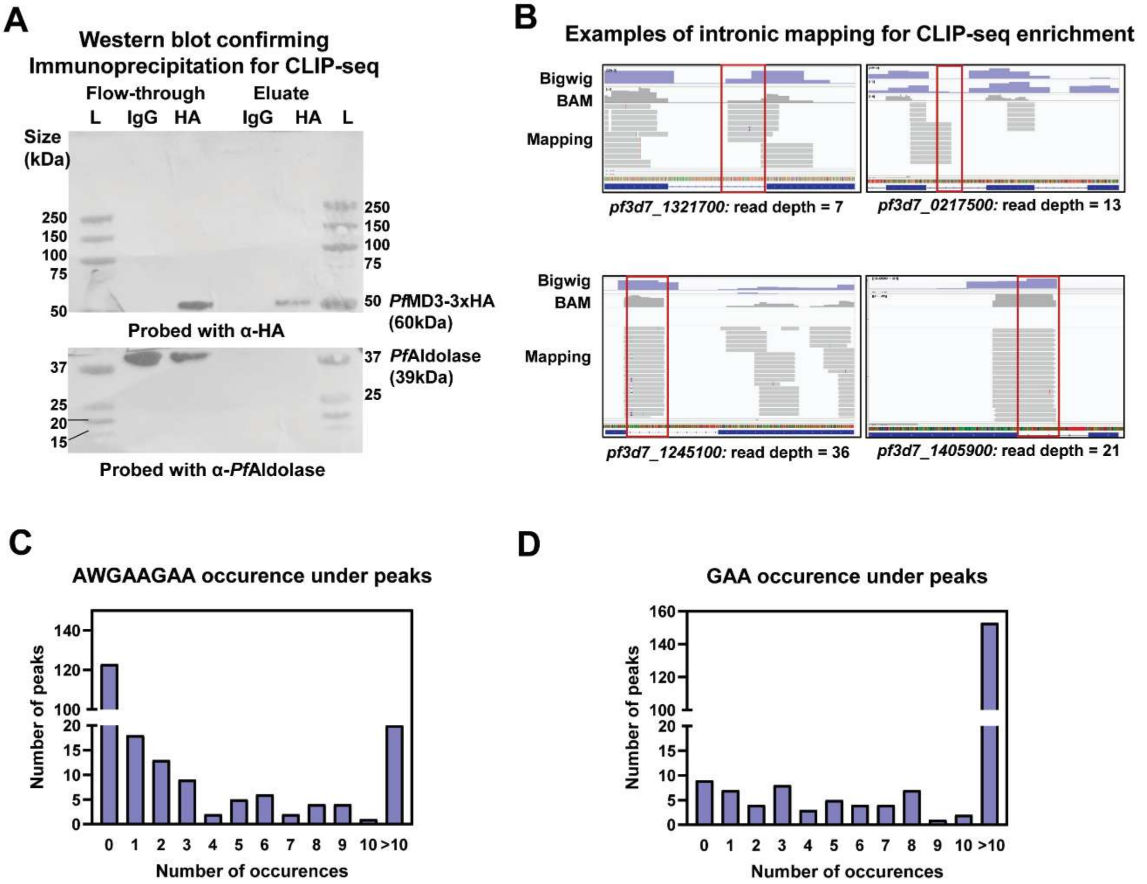


**Figure S5. Confirmation of immunoprecipitation, intronic binding and sequence specificity from *in vivo* CLIP-seq and *in vitro* RBNS results, related to Figure 4.** (**A**) Western blot shows chemiluminescent visualization of anti-HA and anti-*Pf*aldolase staining for the *Pf*MD3-3xHA immunoprecipitation with rat anti-HA antibody (1:2500) and rat anti-IgG antibody (1:2500) respectively. L = ladder, IP = Immunoprecipitation. (**B**) Examples of mapping of enriched CLIP regions to introns in bound transcripts in the IP are shown compared to the input control. Bigwig tracks show input subtracted immunoprecipitation samples while the relative coverage over the introns is shown for BAM files filtered for mapping to unique sites in the genome. Enriched regions were visualized in IGV (v2.1.3). (**C, D**) Number of instances of occurrences of enriched motif from CLIP-seq (AWGAAGAA) or GAA are shown for areas occurring under CLIP-peaks.

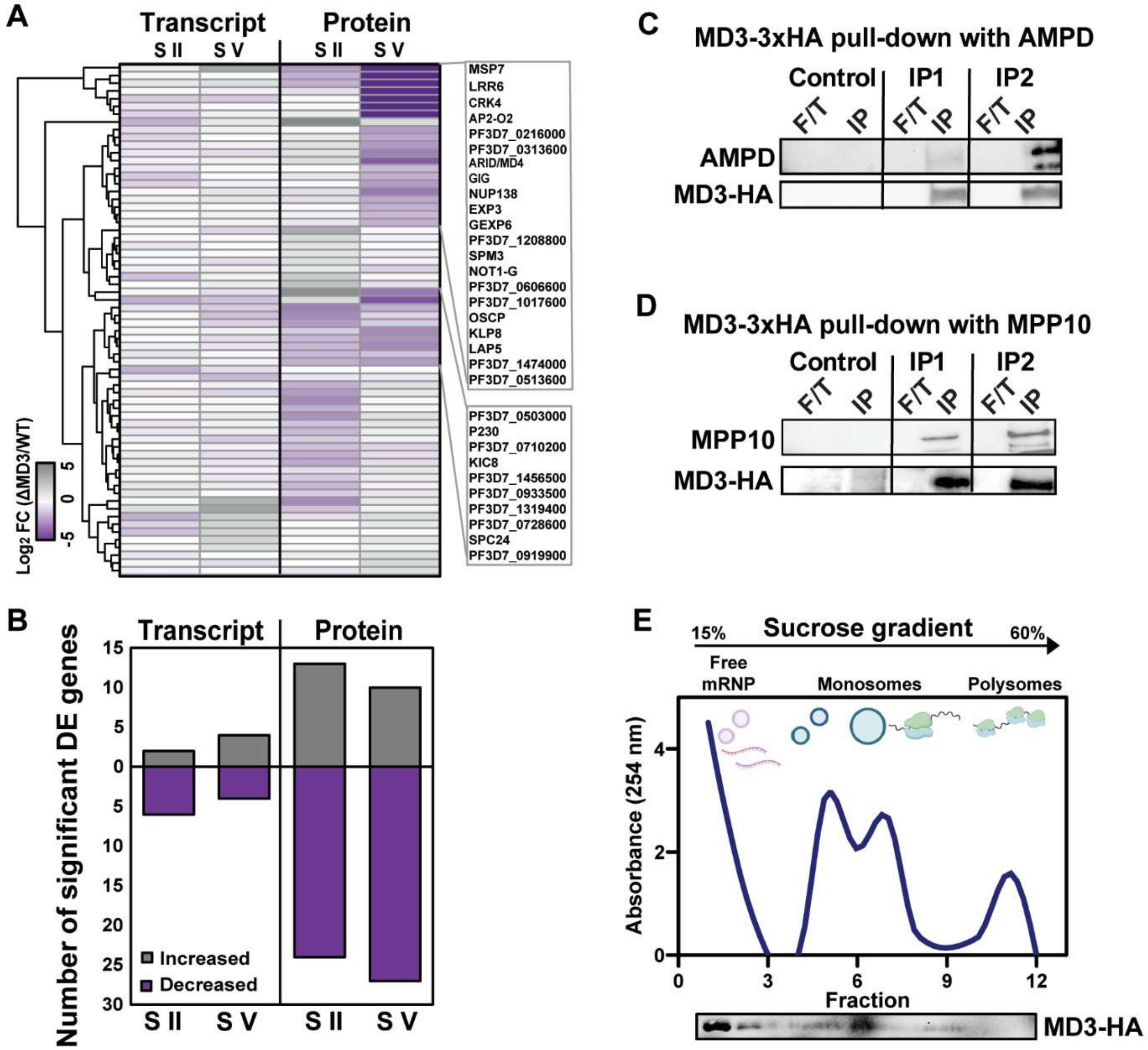


**Figure S6. Gene expression of *Pf*MD3 targets change in Δ*Pf*MD3 parasites, related to Figure 5 and western blot images related to Figure 6.** (**A**) Heatmap representation of $Log_2$ Fold Change of the transcript and protein abundance for the mRNA targets of *Pf*MD3 at Stage II and V of Δ*Pf*MD3$^{1.1}$ gametocyte development. (**B**) The histogram indicates the number of *Pf*MD3 mRNA targets detected by CLIP-seq that are differentially abundant on the protein or transcript level for each stage respectively. Cropped western blot shows confirmation of interaction of *Pf*MD3-3xHA with recombinant 6xHis-tagged MPP10 (**C**) and AMP deaminase (**D**) using protein pull down from *Pf*MD3-3xHA tagged lysate for n = 2 biological replicates (F/T = flow through, IP = immunoprecipitated sample). (**E**) Shows a western blot image and A254 trace related to sucrose gradient (fractions 1-11) profiling obtained for MD3-3xHA from a second biological replicate obtained as in Figure 6E.
